# Fibroblast Activation Protein Defines an Aggressive, Immunosuppressive EMT-associated Tumor Subtype in Highly Inflamed Localized Clear Cell Renal Cell Carcinoma

**DOI:** 10.1101/2024.10.27.620479

**Authors:** Teijo Pellinen, Lassi Luomala, Kalle E Mattila, Annabrita Hemmes, Katja Välimäki, Oscar Brück, Lassi Paavolainen, Elisa Kankkunen, Harry Nisén, Petrus Järvinen, Leticia Castillon, Sakari Vanharanta, Paula Vainio, Olli Kallioniemi, Panu M. Jaakkola, Tuomas Mirtti

## Abstract

Clear cell renal cell carcinoma (ccRCC) is among the most immune-infiltrated cancers, where high inflammation correlates with poor clinical outcomes. While epithelial-to-mesenchymal transition (EMT), immunosuppression, and cancer-associated fibroblasts (CAFs) have been linked with poor prognosis in metastatic ccRCC, their relevance and interplay in localized disease remain uncertain. To address this gap, we performed single-cell spatial profiling of highly inflamed localized tumors to characterize interactions between EMT-associated cancer cells and immune subsets, aiming to identify tumor microenvironment cell subsets and states responsible for the poor prognosis in highly inflamed ccRCC. Multiplexed immunofluorescence imaging with a 33-marker panel was applied to 1,728 tissue cores collected from tumor centers, invasive borders, and adjacent benign tissue from 435 localized ccRCC patients. Independent discovery (n=196) and validation (n=239) cohorts were used to confirm the significance of identified prognostic markers. Hierarchical clustering revealed TME subsets enriched for CD45⁺ immune cells, CD31⁺ endothelial cells, or fibroblasts. High CD45⁺ cell density significantly correlated with poor recurrence-free survival across tumor regions, including tumor center, invasive border, and adjacent benign tissue. Within tumors exhibiting high CD45⁺ infiltration, EMT-associated cancer cells strongly expressed fibroblast activation protein (FAP) at invasive borders, independently predicting worse prognosis and increased risk of liver metastasis. Furthermore, CD45^high^ tumors with elevated FAP expression featured immunosuppressive FAP⁺ CAFs, M2-like macrophages, exhausted T cells, and regulatory T cells. Tumor-cell-specific FAP expression was validated as an independent prognostic biomarker in both early-stage localized ccRCC (pT1-2) and patients subsequently developing metastases who received sunitinib, highlighting its potential for improved patient stratification.

**Significance:** FAP expression identifies an aggressive, immunosuppressive EMT-associated subtype of localized ccRCC, uncovering critical interactions between cancer cells and the TME and revealing novel biological insights and therapeutic vulnerabilities driving early-stage ccRCC progression.

**Graphical Abstract:** *Tumor-cell FAP Expression Defines an Aggressive, Immunosuppressive EMT Subtype in Highly Inflamed Localized ccRCC:* 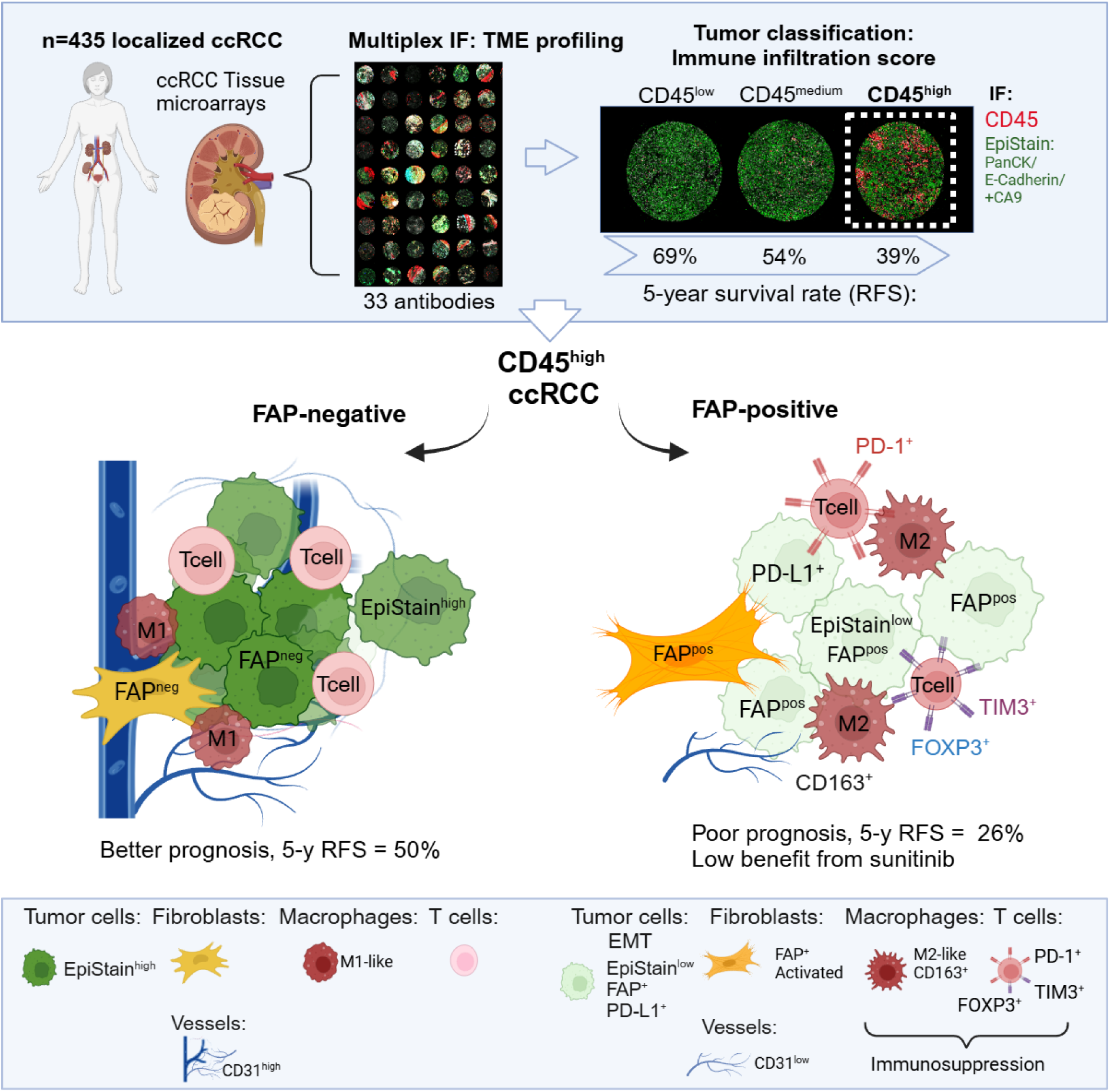 Localized ccRCC tumors were stratified by CD45⁺ immune infiltration, revealing that CD45^high^ tumors were associated with significantly poorer recurrence-free survival (5-year RFS rate: 39%). Among these, tumor cell-specific expression of fibroblast activation protein (FAP) identified a particularly aggressive subtype marked by epithelial-to-mesenchymal transition (EMT), reduced endothelial content, and an immunosuppressive tumor microenvironment enriched with M2-like macrophages, regulatory T cells, exhausted T cells, and FAP⁺ cancer-associated fibroblasts (CAFs). FAP⁺ tumors had an even lower 5-year RFS rate of 26%, compared to 50% in FAP⁻ CD45^high^ tumors. Tumor-cell FAP thus provides an additional layer of risk stratification within the CD45^high^ subgroup and reflects a biologically distinct, immune-evasive tumor state. These findings highlight FAP as both a potential therapeutic target and imaging biomarker in localized ccRCC.

## INTRODUCTION

Clear cell renal cell carcinoma (ccRCC) accounts for over 75% of all renal cell carcinomas (RCCs) and is associated with poor prognosis (1). At diagnosis, 8–19% of patients present with metastases, resulting in 5-year survival rates of only 9–27% (1,2). Even after surgical treatment for localized tumors, up to 40% of patients eventually develop metastases and are likely to die of RCC (3). Since the early 2000s, the use of antiangiogenic tyrosine kinase inhibitors (TKIs), immune checkpoint blockade (ICB) therapy, and their combinations has significantly prolonged survival in patients with metastatic RCC (4,5). Additionally, adjuvant ICB with pembrolizumab has improved disease-free and overall survival in patients with localized ccRCC at intermediate-high and high risk of recurrence after nephrectomy (6,7). However, the optimal adjuvant treatment for localized ccRCC remains unclear, as most patients do not benefit from neoadjuvant or adjuvant ICB in contemporary clinical trials (8–10). Furthermore, despite extensive research, novel liquid or tissue biomarkers have not yet been adopted into routine clinical practice to guide treatment decisions in the adjuvant and metastatic settings (5). Thus, there is an unmet need for personalized treatment strategies for patients with ccRCC.

ccRCC is among the most vascularized and immune-rich cancer types (11). Previous research suggests that patients with ccRCC exhibiting gene expression signatures indicative of high angiogenesis tend to have improved survival outcomes, applicable to both treatment-naïve individuals and those receiving TKIs (11–17). In contrast to most other solid tumors, increased CD8⁺ cell densities and immune cell signatures have been linked to poor outcomes in metastatic ccRCC (12,15,18). However, the association between immune cell signatures and disease recurrence in localized ccRCC is less conclusive (17). Prior studies have also identified elevated levels of immunosuppressive exhausted T cells and M2-like macrophage subsets associated with disease progression (19–22) and response to TKIs and ICB (12,13,23,24). However, most immune cell profiling studies with sufficient sample sizes for survival analysis have utilized bulk RNA sequencing. This method can lead to significant misinterpretations due to mixed sampling of cancerous and normal areas in complex tumors like ccRCC (24,25). In such tumors, understanding the spatial distribution of single cells—such as differences between the tumor center, invasive edge, and tumor-adjacent normal cells—is crucial for accurately deciphering the dynamics of cancer progression in relation to its microenvironment (26).

Recent studies examining the invading tumor edge in ccRCC have observed specific spatial associations between tumor cells undergoing epithelial-mesenchymal transition (EMT) and tumor-associated macrophages (TAMs), T cells, and specific cancer-associated fibroblasts (CAFs) (26–28). Davidson et al. (28) identified a subset of CAFs expressing myofibroblast markers (e.g., αSMA, FAP, FN1) that enrich at the invading tumor edge and correlate with an EMT^high^ phenotype. These observations align with a recent RNA-seq-based molecular classification of advanced RCC into seven distinct subtypes (15). This classification identified a stromal cluster (cluster 6), characterized by genes associated with’activated’ fibroblasts, such as FAP (fibroblast activation protein), FN1 (fibronectin), POSTN (periostin), and various collagens. Notably, this cluster was linked to the poorest survival regardless of the treatment regimen in metastatic RCC (15). Although these studies did not directly link single-cell spatial data with survival outcomes, they highlighted transcriptomic associations between stromal cells—especially FAP⁺ subsets—EMT, and poor outcomes in ccRCC (15,28). Validating these findings in localized tumors at the protein level remains essential, especially considering the spatial context within the tumor microenvironment.

The immune landscape of ccRCC demonstrates extensive heterogeneity within and between patients (13). Emerging evidence indicates that high immune infiltration—particularly of lymphoid and myeloid subsets—is associated with differential benefits from immune checkpoint blockade (ICB) therapy: generally improved progression-free survival (PFS) for lymphoid-rich tumors and worse PFS for myeloid-rich tumors (13,29).

However, the predictive value of these immune cell infiltrations—and their associations with tumor cells and other TME cells—for the progression of localized disease remains unclear. A deeper understanding of spatial cellular profiles within the TME is critical for informing the design of future immuno-oncology clinical trials and developing personalized therapeutic strategies for patients with localized ccRCC. To address this, we utilized multiplexed immunofluorescence (mIF) to analyze 1,728 tissue microarray (TMA) cores from localized ccRCC tumors, employing multi-focal sampling from the tumor center, invasive edge, and tumor-adjacent normal areas.

We spatially characterized the interplay of specific tumor cell states with immune cell subsets and stromal cancer-associated fibroblasts (CAFs) in ccRCCs with high leukocyte infiltration and studied their associations with disease recurrence.

## MATERIALS AND METHODS

### Ethical approvals and data handling

This study was approved by the Institutional Review Boards of Helsinki University Hospital (Ethical Committee of Helsinki University Hospital, diary number HUS/1040/2018) and Turku University Hospital (License number T06/032/15). Informed consent was waived in accordance with Finnish legislation on the secondary use of health data. All data were anonymized before statistical analysis to comply with data protection regulations.

### Patient cohorts and tissue microarrays

Formalin-fixed, paraffin-embedded (FFPE) surgical specimens from 435 treatment-naïve patients with localized (N0M0) ccRCC who underwent partial or radical nephrectomy at Helsinki (n=196) and Turku (n=239) University Hospitals between 2003 and 2013 were utilized in this study. Patients with distant metastases (M1), regional lymph node metastases (N1), a history of kidney cancer, or multiple kidney tumors at diagnosis were excluded. These specimens and the corresponding tissue microarrays (TMAs) had been previously collected and constructed, as described in (30). The TMAs included two cores from the central tumor (1.0 mm for the Helsinki cohort, 1.5 mm for the Turku cohort), two from the tumor border, and two from adjacent benign kidney tissue.

Digital slides were scanned and annotated for TMA construction.

In the Helsinki cohort, patient selection was based on recurrence-free survival (RFS) times: 70 patients with the shortest RFS and 150 with the longest RFS were chosen for TMA construction. After excluding non-clear cell histology cases, 196 patients with localized ccRCC were included. Two pathologists selected representative tissue blocks. Clinical and pathological data—including TNM stage, tumor grade (4-tier Fuhrman/ISUP, depending on the time of diagnosis), sarcomatoid differentiation, necrosis, and survival outcomes—were collected from medical records. The follow-up cutoff date was September 9, 2019.

The Turku cohort included 239 patients treated with nephrectomy between 2005 and 2014, representing a continuous, population-based series. The same clinicopathological inclusion criteria were applied as in the Helsinki cohort. Clinical and pathological features, including RFS and disease outcomes, were extracted from electronic medical records. Follow-up continued until July 11, 2019. The patient characteristics are shown in **Supplementary Table S1**.

### Multiplexed immunofluorescence staining and imaging

Our in-house multiplexed immunofluorescence (mIF) protocol involves multiple rounds of staining and scanning. The first two antibodies in each panel are amplified via a tyramide signal amplification (TSA) reaction and detected using Alexa Fluor 488 (AF488) or Alexa Fluor 555 (AF555). Subsequent antibodies are stained and detected with AF647 and AF750-conjugated secondary antibodies. After each staining round, slides are scanned, and antibodies are removed through bleaching and denaturation. The antibody panels, along with antibody details and secondary reagents, are shown in **Supplementary Table S2**. The detailed mIF protocol will be deposited to https://www.protocols.io upon acceptance of the manuscript.

TMA sections (3.5 μm) were deparaffinized and rehydrated, followed by heat-induced epitope retrieval (HIER) in Tris-EDTA buffer (pH 9) at 99°C for 20 minutes using the PT module (Epredia, USA). Endogenous peroxidase activity was blocked with 0.9% hydrogen peroxide (H₂O₂) in Tris-buffered saline (TBS) for 15 minutes at room temperature (RT). Tissue sections were then blocked with 10% goat serum in TBS with 0.05% Tween-20 (TBST) for 15 minutes at RT before antibody application.

The first two primary antibodies were incubated for 1 hour at RT. Detection was performed using anti-mouse or anti-rabbit poly-HRP (1:5 in TBST; BrightVision, Immunologic, the Netherlands), followed by incubation with tyramide-AF488 (1:100; Thermo Fisher Scientific, USA) in TBST with 0.0015% H₂O₂ for 15 minutes at RT. Residual HRP activity was quenched with 0.9% H₂O₂ treatment. The second antibody was detected using tyramide-AF555 (Thermo Fisher Scientific, USA) following the same procedure. After HIER and blocking, the third and fourth antibodies were applied and detected using secondary antibodies conjugated to AF647 and AF750 (1:300) along with DAPI (1.6 μg/ml). Slides were washed and mounted using ProLong Gold Antifade Reagent (Thermo Fisher Scientific, USA) and scanned with a Zeiss Axioscan.Z1 slide scanner (Carl Zeiss, Germany) using the Colibri7 light source and filter set 112 (wavelengths 365 nm [DAPI], 488 nm, 555 nm, 647 nm, and 750 nm). Following scanning, coverslips and fluorochromes were removed by immersing slides in TBST, followed by bleaching with 24 mM NaOH and 4.5% H₂O₂. After HIER, additional antibody pairs were applied and detected with AF647 and AF750. For CAF panel 1, both primary antibodies were rabbit-derived and detected sequentially using tyramide reagents, with HIER between applications. The FAP antibody was detected via a tyramide-biotin reaction with streptavidin-AF750. For CAF panel 2, the first antibody was goat-derived and detected with anti-goat HRP and tyramide-AF488, followed by tyramide-AF555 for the second antibody.

FAP antibody was validated using CRISPR-Cas9 knockout in WPMY-1 cell line. The knockout of the FAP gene in the WPMY-1 cell line was achieved using lentivirus-mediated gene delivery (31). Lentiviral particles containing FAP_RNA1 and FAP_RNA2 constructs were produced in HEK293FT cells utilizing a three-plasmid system (psPAX2, pCMV VSVG, and a FAP-expressing vector). After collecting the viral supernatants, WPMY-1 cells were infected with the FAP_RNA1 or FAP_RNA2 virus in the presence of polybrene (8 µg/ml) to enhance infection efficiency. Post-infection, the cells were selected using puromycin-containing media (1 µg/ml), ensuring that only successfully transduced cells survived. Following infection, single-cell cloning was performed using serial dilution in 96-well plates. This technique allowed the isolation of single-cell clones, and three independent clones were identified as fully FAP knockout, confirmed by immunohistochemical staining (**Supplementary Fig. S4B**).

### Image processing and feature extraction

Zeiss image data were exported as single-channel grayscale TIFFs at 25% and 100% resolution. Tissue cores were annotated and cropped using FIJI/ImageJ using the DAPI channels, and image coordinate-based cropping was applied to other channels using MATLAB (version 2023a; MathWorks Inc., Natick, MA, USA). Image registration using DAPI channels as references between different image rounds was performed with MATLAB, as described previously (32). Low-resolution images were used to create RGB images for segmentation in ilastik (version 1.3.3) (33), focusing on red blood cell autofluorescence using the DAPI and TSA-488 channels. In the TME panel, ilastik was also applied to segment epithelial regions (markers: CA9, PanCk, E-cadherin), generating a binary epithelial mask (EpiMask). In CAF panel 1, ilastik was used to differentiate stromal areas from epithelial areas using RGB input images composed of PDGFRB and αSMA channels (stroma, green), the panEpi channel (blue), and DAPI (red). In the T cell, macrophage, and CAF Panel 2, ilastik was not used for segmenting epithelium/stroma due to inadequate coverage for training.

Full-resolution DAPI images were used for nuclear segmentation using the nucleAIzer deep-learning algorithm (34). Cells were defined by dilating the nuclei by 3 pixels, and cell features—including marker intensities and binary masks—were extracted using CellProfiler (version 4.2.1) (35). Cells in contact with autofluorescent red blood cells were excluded from the analysis.

### Cell classification

All analyses were performed using Jupyter Notebook (Python 3.6.8). Cells were classified as either epithelial (Epi^+^) or non-epithelial (Epi^−^)using different approaches due to varying stromal/epithelial stains in the panels. In the TME panel, Epi^+^ cells were defined as positive for the ilastik machine learning-derived EpiMask (>50% pixel coverage), but negative for immune cells (CD45, CD3, CD20, CD11b) and stromal cells (PDGFRB, D2-40, CD31). In CAF panel 1, stromal cells were positive for the ilastik-derived Stroma mask (>10% pixel coverage) and negative for EpiMask (<90%). In the T cell panel, macrophage panel, and CAF panel 2, panEpi intensity threshold was applied for Epi^+^ classification. Positivity thresholds for each marker intensity were determined by visual inspection.

Distinguishing epithelial from non-epithelial cells with high accuracy can be challenging in mIF staining due to the limited number of markers available. However, the TME panel provided the best epithelial cell coverage in this study. Therefore, epithelial characteristics, such as EpiStain intensity for EMT marker analysis, were measured in the TME panel.

### Quantification of cell populations

TMA cores with poor quality—such as broken tissue or poor focus—were excluded from the analyses. This process was performed separately for each panel, leading to slightly different final numbers of TMA cores and patients across the analyses. For quantification, cell counts and proportions were averaged by taking the mean of the replicate TMA cores. Proportions were calculated based on specific cell categories, such as the proportion from Epi⁻ cells (**Fig. 2B-K; 4A**), the proportion of CD45^+^ cells (**Fig. 5A**), or the proportion of PD1^+^TIM3^+^CD3^+^CD8^+^ cells among all CD3^+^CD8^+^ cells (**Fig. 5C**). In CAF Panel 1, we classified CAFs based on our previously established system from studies in non-small cell lung carcinoma (NSCLC) and cutaneous squamous cell carcinoma (cSCC),(36,37) assigning categories from CAF1 to CAF15, each defined by a unique set of quadruple markers (PDGFRA, PDGFRB, FAP, αSMA). The CAF1 to CAF15 cell subsets were summed, and each CAF subset was divided by the total sum to yield a proportion of the total CAFs. The same quantification approach was applied to CAF Panel 2 (CAF16 to CAF30).

**Figure 1.**
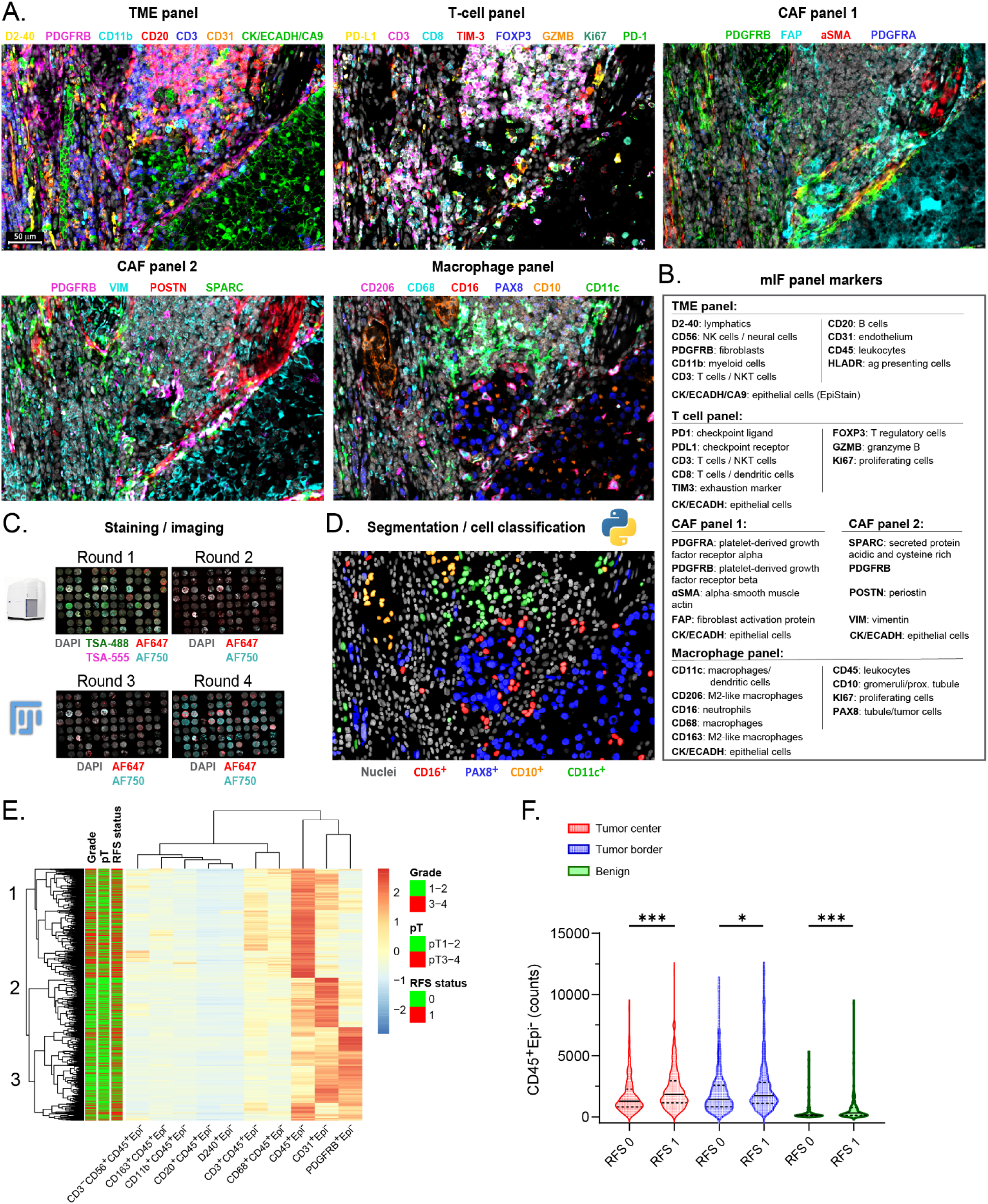
Single-cell multiplexed IF reveals a leukocyte cell cluster associated with disease progression in localized ccRCC. **(A)** Representative multi-channel images from multiplexed immunofluorescence (mIF) antibody panels, depicting consecutive sections from the same ccRCC tissue sample focused on a tumor border area. Scale bar represents 50 μm. **(B)** mIF antibody panels with markers listed according to their primary targets for immune panels and as protein names for CAF panels. **(C)** Illustration of the mIF protocol, including four staining and imaging cycles with Tyramide-signal amplification (TSA) in Round 1 (channels 488/555). Cores were annotated, cropped using ImageJ/FIJI, and registered in MATLAB using DAPI as a reference. AF, Alexa Fluor dye. **(D)** Illustration of single-cell segmentation (nucleAIzer(34)) and cell classification based on mean intensity thresholding, exemplified using the macrophage panel image. **(E)** Heatmap displaying Euclidean clustering (rows and columns) of absolute counts for major cell subsets in individual TMA cores (n = 1728). Annotations include Fuhrman grade, pathological stage (pT), and recurrence-free survival status (RFS 0 for non-recurrent, RFS 1 for recurrent with event: metastasis or death). **(F)** Violin plots illustrating the distribution differences of CD45^+^Epi⁻ cell densities between non-recurrent (RFS 0) and recurrent (RFS 1) patient groups. Data are shown for tumor center (n = 405), tumor border (n = 391), and benign areas (n = 355). The lines represent medians and interquartile ranges (IQR). ***, p < 0.001; *, p < 0.05 (Mann-Whitney u test).

**Figure 2.**
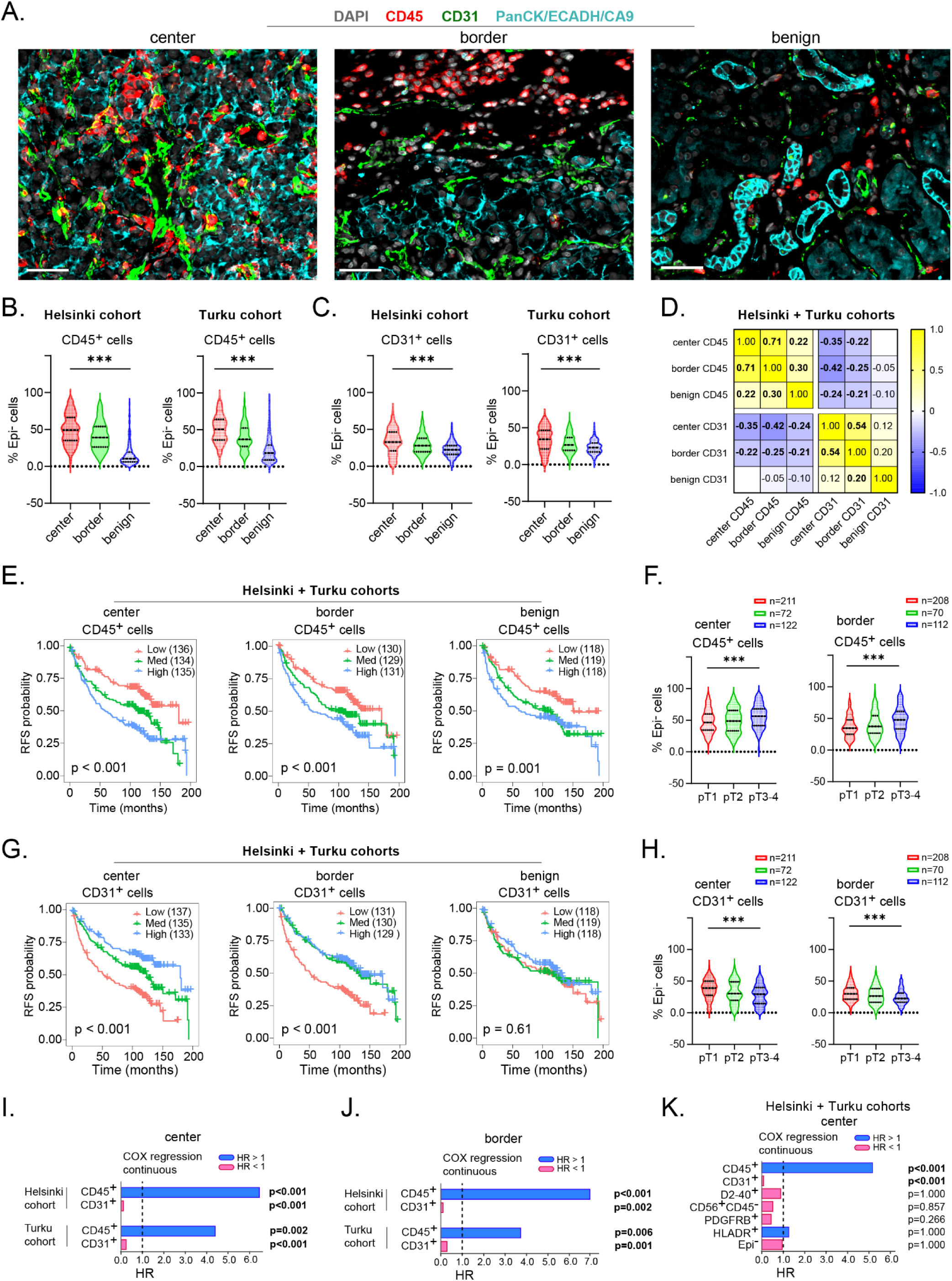
Leukocyte and endothelial cell densities determine RFS in localized ccRCC. **(A)** Example mIF images showing CD45 and CD31 staining alongside kidney epithelium stain (EpiStain) containing Pan-cytokeratin (PanCK), E-cadherin (ECADH), and carbonic anhydrase IX (CA9). Scale bar, 50 µm. **(B)** Relative densities of CD45^+^Epi⁻ cells as a proportion of total Epi⁻ cells in tumor center, border, and benign areas in Helsinki (n = 178) and Turku (n = 227) localized ccRCC cohorts. ***, p < 0.001 (Kruskal-Wallis test). **(C)** Similar to (B), but for CD31^+^Epi⁻ cells. **(D)** Correlation matrix (Pearson r) showing the relationship of CD45^+^Epi⁻ and CD31^+^Epi⁻ cell densities (as a proportion of total Epi⁻ cells) across tumor areas. Bold values indicate p < 0.05. **(E)** Kaplan-Meier survival curves for recurrence-free survival (RFS) based on trichotomized proportions of CD45^+^Epi⁻ cells in different tumor areas. The categorization into low, medium, and high counts is based on a three-tier equal categorization derived separately from the Helsinki and Turku ccRCC cohorts before cohort merging. This explains the slightly different n values for each category. p-values from log-rank test. **(F)** Density distributions of CD45^+^Epi⁻ cells in relation to pT staging groups in tumor center and border areas. ***, p < 0.001 (Kruskal-Wallis test). **(G)** Similar to (E), but for trichotomized proportions of CD31^+^Epi⁻ cells. p-values from log-rank test. **(H)** Similar to (F), but for CD31^+^Epi⁻ cells. ***, p < 0.001 (Kruskal-Wallis test). **(I-K)** Univariate Cox regression survival analysis using continuous values for the indicated cell subset proportions (all subsets are fractions of Epi⁻ cells, except for Epi⁻, which is a fraction of total cells) in tumor center (I) and border (J) areas, and in the merged cohort for tumor center areas (K). HR represents hazard ratio.

### Visual scoring of mesenchymal markers in tumor cells

Visual inspection of the multiplexed immunofluorescence (mIF) images of the stained tissue microarrays (TMAs) suggested that automated digital analysis of mesenchymal markers within the epithelial compartment was problematic. This difficulty arose because the detection and segmentation of epithelial and stromal cells were not optimal in panels other than the tumor microenvironment (TME) panel, which included a higher number of epithelial antibodies such as carbonic anhydrase IX (CA9) to achieve better epithelial cell coverage.

To address this issue, visual scoring of mesenchymal markers—including platelet-derived growth factor receptor alpha (PDGFRA), platelet-derived growth factor receptor beta (PDGFRB), fibroblast activation protein (FAP), alpha-smooth muscle actin (αSMA), secreted protein acidic and rich in cysteine (SPARC), vimentin (VIM), periostin (POSTN), and programmed death-ligand 1 (PD-L1)—was performed by two scientists (T.M. and T.P.). In cases of disagreement, a consensus score was reached after re-evaluation. Hematoxylin and eosin (H&E), PAX8, and EpiStain (from the TME panel) stained samples from consecutive or near-consecutive histological sections were used as reference images to guide accurate identification of epithelial regions.

Among these markers, PDGFRA, PDGFRB, αSMA, and POSTN were negative in epithelial cells. In contrast, FAP, SPARC, VIM, and PD-L1 showed clear positivity in ccRCC cells. FAP and PD-L1 were scored in each tumor TMA core (two from the tumor center and two from the tumor border) as negative (score = 0), weak (score = 1), or strong (score = 2). In Fig. 3M–3P and Fig. S4B–S4E, a cumulative visual scoring was applied for FAP and PD-L1 by summing the scores from all tumor cores for the same patient. For example, cumulative FAP scoring was defined as follows:

**Figure 3.**
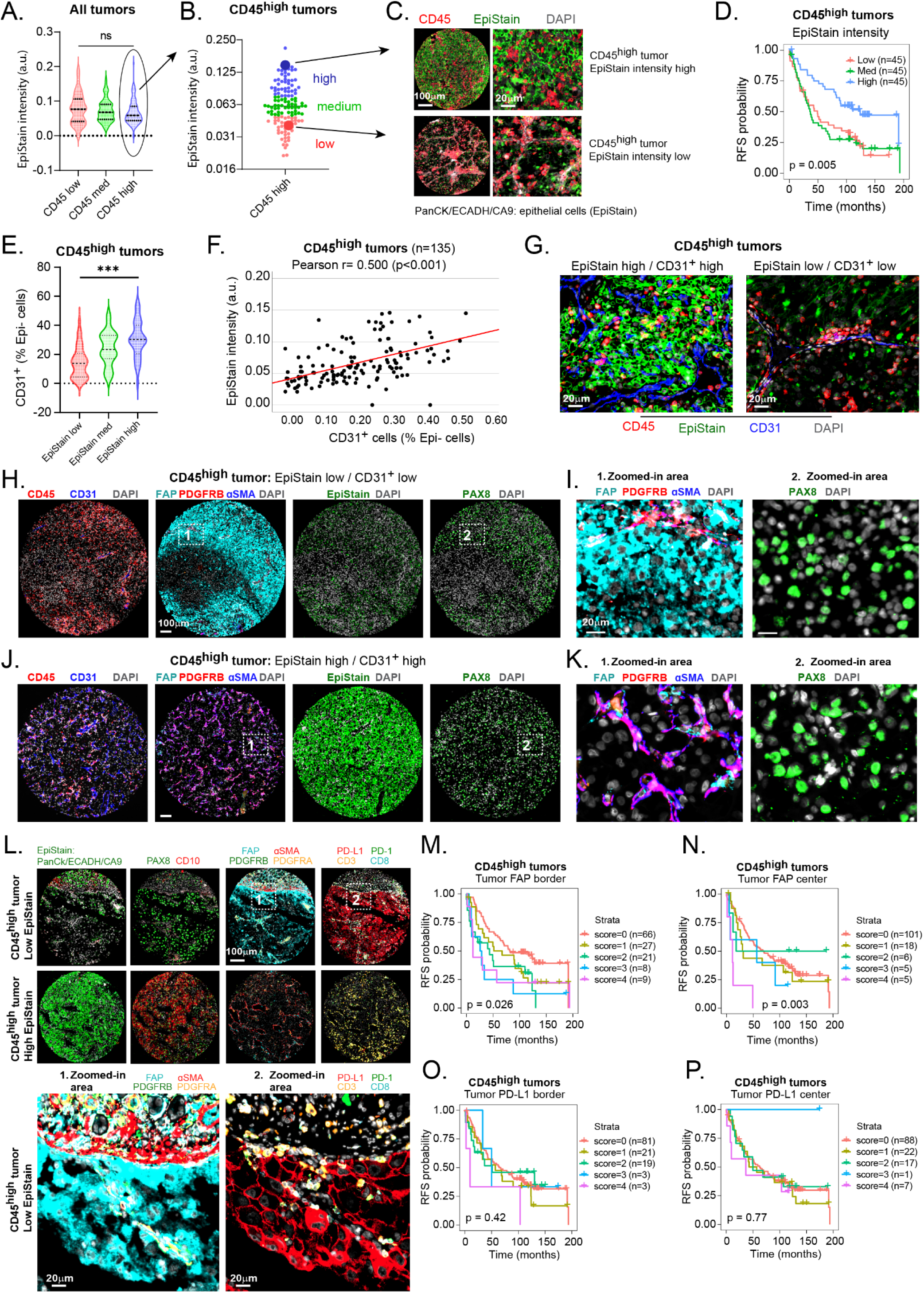
High cancer cell FAP expression identifies aggressive CD45^high^ tumors with EMT and PD-L1 upregulation. **(A)** EpiStain intensity distribution in tumor center cores grouped by low (n=135), medium (n=135), and high (n=135) CD45^+^ cell density. n(ccRCC tumors) = 405, p = 0.098; Kruskal-Wallis test. **(B and C)** Classification of CD45^high^ tumors (n = 135) into equal-sized low, medium, and high EpiStain intensity groups, with representative mIF for EpiStain^high^ and Epistain^low^ cases. **(D)** Kaplan-Meier survival analysis for CD45^high^ tumor patients (n = 135), categorized by EpiStain intensity. p-values from log-rank test. **(E-G)** Violin plots (E), scatter plots (F), and example mIF images (G) showing the correlation between EpiStain intensity and CD31^+^ cell density in tumor center of CD45^high^ ccRCCs. **(H-K)** mIF images comparing CD45^high^ ccRCCs with contrasting phenotypes. (H) EpiStain^low^ / CD31^+^low phenotype with tumor-specific FAP expression (I, zoomed view); (J) EpiStain^high^ / CD31^+^high phenotype with focus area shown in (K). **(L)** Representative mIF images of CD45^high^ ccRCCs with low (upper panel) and high (lower panel) EpiStain-associated phenotypes. **(M-N)** Kaplan-Meier curves for tumor cell-specific FAP expression in CD45^high^ ccRCCs (n = 135) and All ccRCCs (n = 405) based on 3-tier visual scoring. FAP was scored (negative = 0, weak = 1, strong = 2) in tumor border (n = 2) and center TMA cores (n = 2) separately and the scores were summed, yielding final cumulative scores from 0 to 4 (see Methods for details). p-values from log-rank test. **(O-P)** Similar analysis as in (M-N), but for tumor cell-specific PD-L1 expression.

Score = 0: All cores negative for tumor FAP.

Score = 1: One core weakly positive for tumor FAP.

Score = 2: Two cores weakly positive or one core strongly positive for tumor FAP. Score = 3: Two cores weakly positive and one core strongly positive for tumor FAP. Score = 4: Two cores strongly positive for tumor FAP.

In **Tables 1, S3, S5**, and **Supplementary Fig. S5**, any positive expression (scores = 1 or 2) in either replicate core from the tumor border or center (analyzed separately or combined) was considered a positive score for a marker. In **Tables S4, S6, and S7**, as well as in **Fig. 4C, 5D–5E, 6, S6**, and **S8G–S8H**, the maximum score of tumor FAP or tumor PD-L1 observed in any tumor core (two from the center and two from the border) was recorded (negative = 0, weak = 1, strong = 2).

**Figure 4.**
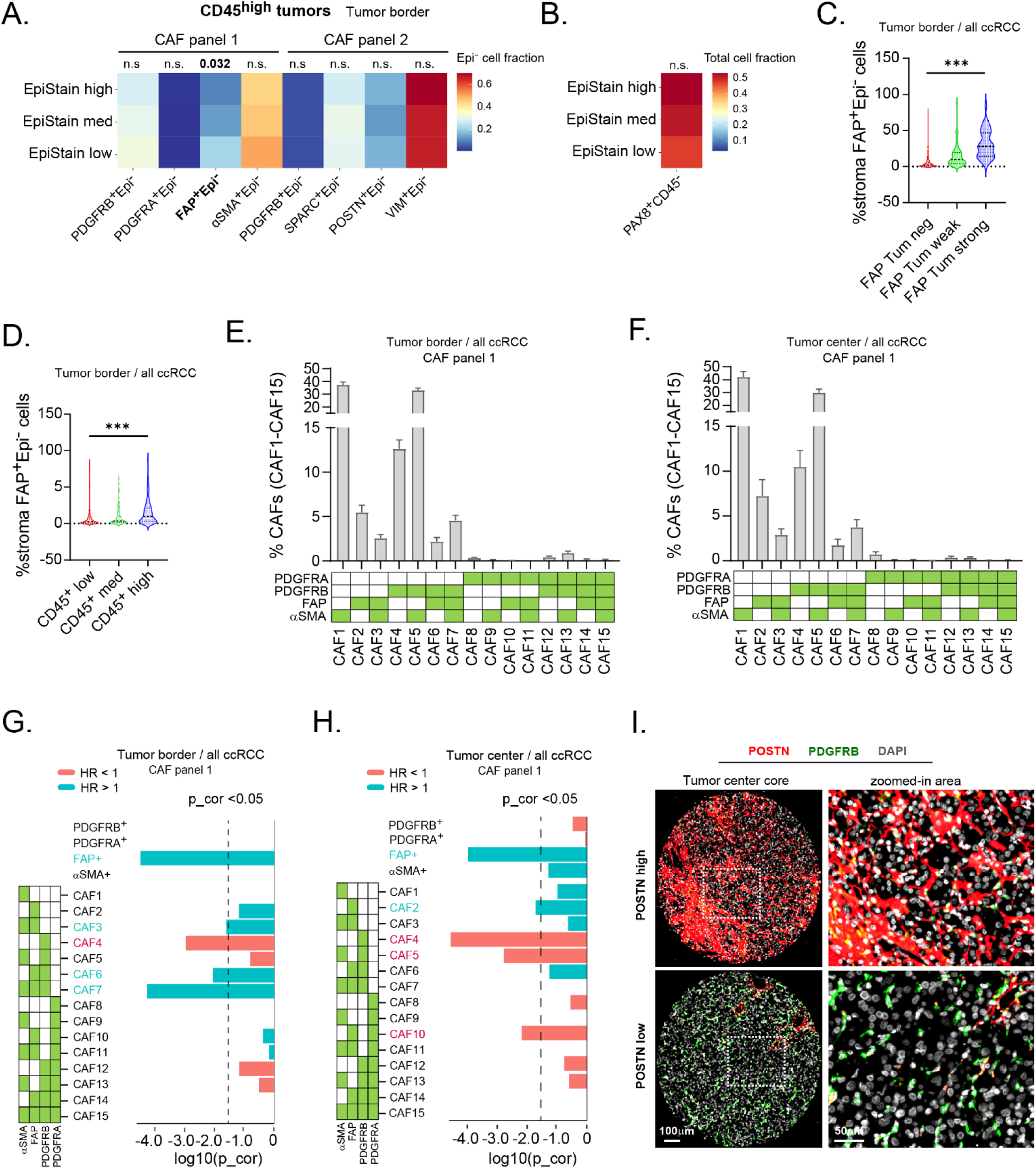
FAP^+^ cancer-associated fibroblasts are enriched at tumor edges in CD45^high^, EMT^+^ ccRCC. **(A)** Distribution of single marker defined cell subsets in tumor border area across EpiStain classes in CD45^high^ ccRCCs (n = 135). The colored values represent cell proportions from total stroma (Epi⁻) cells. Kruskal-Wallis test. **(B)** PAX8^+^CD45^-^ cell subset distribution at tumor border across EpiStain classes in CD45^high^ ccRCC tumors. The colored values represent cell proportions from total cells. Kruskal-Wallis test. **(C)** FAP^+^ stromal cell distribution across tumor FAP classified ccRCCs (n = 394) ***p < 0.001; Kruskal-Wallis test. **(D)** Tumor border FAP^+^ stroma cell distribution across CD45 classified ccRCCs (n = 394) ***p < 0.001; Kruskal-Wallis test. **(E and F)** Relative distribution of CAF panel 1 multi-marker defined stromal cell subsets in tumor border (E) (n = 394) and center (F) (n = 414). Each CAF subset abundance is relative to the sum of the 15 CAFs as in (37). Bars represent mean values and error bars 95% confidence interval. **(G and H)** Univariate Cox regression survival analysis using continuous values for the indicated cell subset proportions (all subsets Epi⁻) in tumor border (G) and center (H) areas. Green box represents positivity. HR, Hazard ratio; p_cor, Bonferroni corrected p-value. **(I)** Representative mIF images of ccRCCs from tumor center areas with high density of POSTN^+^Epi⁻ cells (n = 2157; upper panel) and low density of POSTN^+^Epi⁻ cells (n = 133; lower panel).

**Figure 5.**
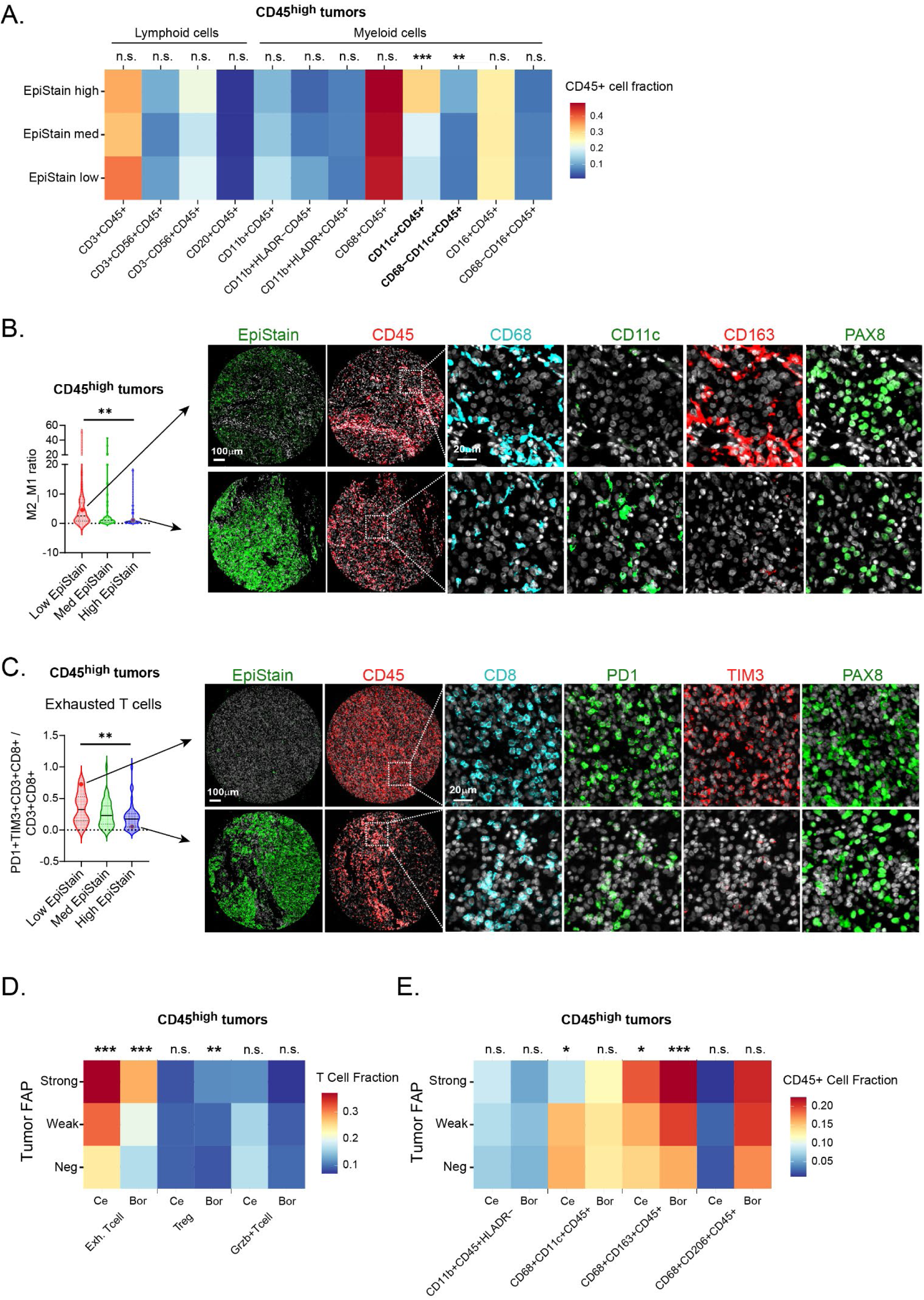
Differential distribution of immune cell subsets in CD45^high^, EMT^high^ ccRCC. (A) Heatmap of immune cell subsets fractions within tumor center cores of CD45^high^ ccRCCs (n = 135), categorized by EpiStain levels. Colors represent the relative fractions of CD45^+^ cells using a reversed RdYlBlu color palette. n.s, not significant; **p < 0.01, ***p < 0.001; Kruskal-Wallis test. (B) Violin plots of M2-to-M1 macrophage ratio categorized by EpiStain levels in CD45^high^ cases (n = 135). The ratio was derived by CD68^+^CD163^+^CD45^+^/CD68^+^CD11c^+^CD45^+^ counts. Tumor center cores. **p < 0.01; Kruskal-Wallis test. Example mIF images illustrate a low and high EpiStain case with associated macrophage phenotypes. (C) Violin plots of exhausted T cells (PD1^+^TIM3^+^CD3^+^CD8^+^/CD3^+^CD8^+^) as relative fractions of total effector T cells (CD3^+^CD8^+^), categorized by EpiStain levels. **p < 0.01; Kruskal-Wallis test. (D) Heatmap of T cell fraction by cell type and tumor FAP level in CD45^high^ cases. Colors represent the relative fractions of effector T cells (CD3^+^CD8^+^) using a reversed RdYlBlu color palette. Tumor FAP scored as negative/weak/strong if either tumor center or border showed positivity (maximum level selected from center/border). T cell fractions as continuous values in tumor center (Ce) and border (Bor) TMA cores (average of replicates). n.s, not significant; **p < 0.01, ***p < 0.001; Kruskal-Wallis test. (E) Same as D, but myeloid cell fractions of CD45^+^ cells. n.s, not significant; *p < 0.05, **p < 0.01, ***p < 0.001; Kruskal-Wallis test.

### Analysis of single-cell RNA-sequencing data

We analyzed single-cell RNA sequencing (scRNA-seq) data from (26). Cells annotated as RCC by the original authors were included. Genes expressed in fewer than 10 cells were excluded. Raw counts for the remaining genes were normalized and transformed using library size normalization followed by square root transformation. The data were then imputed using Rmagic (v2.0.3) with k = 15. Scatter plots of the imputed data were generated to explore the correlation between EMT-associated genes and epithelial markers. The analysis was conducted in R (version 4.0.4) using additional packages including tidyverse (v2.0.0) and Seurat (v5.0.1) (38).

### Statistical analyses

Statistical analyses were performed using GraphPad Prism (version 10.3.1), IBM SPSS Statistics (version 29.0.0.0), RStudio (version 2023.12.0) with R (version 4.2.2; apart from scRNA-seq analysis), and JupyterLab (version 3.0.16) with Python (version 3.6.15). In R, data manipulation and visualization were conducted using the tidyverse package suite (version 2.0.0), including dplyr, ggplot2, readr (version 2.1.4), and readxl (version 1.4.3). Survival analyses were performed using the survival package (version 3.5-7), employing the coxph() function for Cox proportional hazards regression and the survfit() function for Kaplan-Meier survival estimates. The survival endpoint was recurrence-free survival (RFS) and was defined as time from surgery to disease recurrence, death, or until end of follow-up. Survival curves were visualized using the ggsurvplot() function from the survminer package (version 0.4.9), based on ggplot2 graphics, with additional themes applied from the ggthemes package (version 5.0.0). The proportional hazards assumption was tested using the cox.zph() function. The Mann-Whitney U test and Kruskal-Wallis test were used for non-parametric comparisons between two or more groups, respectively. Associations between categorical variables were assessed using Fisher’s exact test or the Chi-square test, as appropriate. When multiple comparisons were made, p-values were adjusted using the Bonferroni correction or controlled for a 10% false discovery rate using the Benjamini-Hochberg procedure. Logistic regression analyses were conducted in Python using the statsmodels package (version 0.13.2) to evaluate the likelihood of liver metastasis based on FAP and PD-L1 expression levels, adjusting for clinical covariates. Odds ratios (ORs) and 95% confidence intervals (CIs) were calculated from the regression coefficients.

### Data availability

Data generated in this study will be deposited in Zenodo and made publicly available upon acceptance of the manuscript. Codes used in the analyses will be deposited in GitHub and will be publicly available upon acceptance.

## RESULTS

### Single-cell multiplexed immunofluorescence reveals a leukocyte cluster associated with disease progression in localized ccRCC

We analyzed formalin-fixed, paraffin-embedded (FFPE) surgical specimens from 435 patients with localized (M0N0) clear cell renal cell carcinoma (ccRCC) who underwent radical or partial nephrectomy at Turku and Helsinki University Hospitals. TMAs were constructed using 1.0 to 1.5-mm cores from each resection, sampling areas from the tumor center (n = 2 cores), tumor border (n = 2 cores), and adjacent non-neoplastic cortical kidney tissue (n = 1 core), hereafter referred to as benign tissue. Median postoperative follow-up times were 8.69 years (interquartile range [IQR], (2.08–10.23)) for the Helsinki cohort and 6.01 years (IQR, 2.09–9.86) for the Turku cohort, including registration of metastatic events and recurrence-free survival (RFS) (see clinicopathological variables in **Supplementary Table S1**).

The TMAs were stained using five different 5-to 10-plex antibody panels, comprising a total of 33 antibodies targeting well-established markers of the tumor microenvironment (TME panel), T cells (T cell panel), macrophages/myeloid cells (macrophage panel), and fibroblasts (CAF panels 1 and 2) (**Fig. 1A-B and Supplementary Table S2**). In the TME panel, we utilized an epithelial-specific antibody cocktail, termed EpiStain, which included antibodies against pan-cytokeratin (PanCk), E-cadherin (CDH1), and carbonic anhydrase IX (CA9). This cocktail allowed us to identify a broad spectrum of epithelial (Epi⁺) cells, accommodating varying levels of marker expression across the tumor center, border, and benign areas in the tissue sections. Tissue cores were imaged using fluorescence microscopy and processed to collect marker expression data from spatially segmented individual cell objects (**Fig. 1C**), which were classified as negative or positive for each marker (**Fig. 1D**), as described previously (37).

Hierarchical clustering (Euclidean distance) of cell counts for selected non-epithelial cell subsets across all TMA cores from both cohorts revealed three distinct sample groups (**Fig. 1E**). These groups represented: (1) high immune cell prevalence (CD45⁺), (2) high vascular endothelial cell prevalence (CD31⁺), and (3) high fibroblast prevalence (PDGFRB⁺). Notably, the density of CD45⁺ cells was higher in the primary tumors of patients who developed recurrent ccRCC compared to those who did not experience disease recurrence, as measured across tumor center, border, and benign tissue (Mann-Whitney U test, **Fig. 1F**).

### Leukocyte and endothelial cell densities determine recurrence-free survival in localized ccRCC

Using the tumor microenvironment (TME) multiplex panel, we quantified the relative densities of CD45⁺ leukocytes and CD31⁺ endothelial cells in relation to total non-epithelial cells (Epi⁻) across the tumor center, tumor edge, and adjacent benign tissue (**Fig. 2A**). Both cell types consistently decreased from the tumor center to the border, with a further reduction observed in the adjacent benign areas (Kruskal-Wallis test, p < 0.001; **Fig. 2B and 2C**). CD45⁺ cells and CD31⁺ cells showed an inverse correlation with each other, while both CD45⁺ and CD31⁺ cell densities were positively correlated across different tumor areas (**Fig. 2D**). Higher densities of CD45⁺ cells in the tumor center, border, and adjacent benign areas were associated with shorter recurrence-free survival (RFS) and higher pathological tumor stages (pT) (**Fig. 2E and 2F**). Conversely, higher densities of CD31⁺ cells were associated with longer RFS and lower pT stages (**Fig. 2G and 2H**). Cox regression analysis of CD45⁺ and CD31⁺ cell densities revealed that the significant associations with RFS remained consistent when evaluated as continuous variables in tumor center and border cores separately in the two cohorts (**Fig. 2I and 2J**). The robustness of the multiplexed immunofluorescence (mIF) staining and survival analyses was confirmed by validating the Kaplan-Meier and Cox regression analyses for CD45⁺ cell densities using data from another experiment, the macrophage multiplex panel (**Supplementary Fig. S1**). Other general cell populations, such as lymphatic endothelial cells (D2-40⁺), fibroblasts (PDGFRB⁺), or the total non-epithelial cell population (Epi⁻) identified within the TME multiplex panel showed no significant association with RFS in the combined cohort (**Fig. 2K**). Overall, the mIF profiling of these localized ccRCC cohorts reflected transcriptomic signature findings related to angiogenesis, inflammation, and survival previously observed mostly in patients with advanced ccRCC (11,12,15).

### High cancer cell FAP expression identifies aggressive CD45^high^ tumors with EMT and PD-L1 upregulation

Recent studies using single-cell RNA sequencing (scRNA-seq) and bulk sequencing have highlighted the importance of epithelial-to-mesenchymal transition (EMT) in the progression of ccRCC (17,26,28). The loss of epithelial markers such as E-cadherin and cytokeratins—which are included in our EpiStain antibody cocktail— is considered a hallmark of EMT (39). Additionally, carbonic anhydrase IX (CA9), also part of our EpiStain cocktail, can be lost in ccRCC cells,(28) consistent with our re-analysis of scRNA-seq data (**Supplementary Fig. S2**) (26).

We observed considerable variability in EpiStain intensity within Epi^+^ cells across tumors from different patients. However, EpiStain intensity did not significantly vary among tumors with low (mean intensity 0.078), medium (0.073), and high (0.067) CD45^+^ cell levels (**Fig. 3A**). Given the potential of highly inflamed ccRCCs as targets for immunotherapy(29) and their under-characterization at localized disease stages, we focused on patients exhibiting high CD45^+^ cell infiltration in tumor center cores (CD45^high^). We further stratified these patients (n = 135) into three equally sized groups based on EpiStain levels within Epi^+^ cells: low (n = 45), medium (n = 45), and high (n = 45) (**Fig. 3B and 3C**). This classification was based on EpiStain levels at the tumor center because tumor border areas contain a mixture of tumor and non-tumor Epi^+^ cells. Despite this heterogeneity, we found a strong correlation between EpiStain levels in Epi^+^ cells at the tumor center and border areas (Pearson r = 0.7, p < 0.001; **Supplementary Fig. S3**).

Intriguingly, among patients with CD45^high^ tumors, those exhibiting high EpiStain levels in the tumor center had a longer recurrence-free survival (RFS) compared to CD45^high^ patients with low or medium EpiStain levels (log-rank test, p = 0.005; **Fig. 3D**). Furthermore, in CD45^high^ tumors, the EpiStain intensity within Epi^+^ cells of the tumor center correlated with the abundance of CD31^+^ cells in the stromal compartment (**Fig. 3E–3G**), reflecting their association with favorable RFS.

Next, we investigated whether lower EpiStain levels in tumor cells are associated with increased mesenchymal marker expression, indicative of EMT or partial EMT. Due to potential challenges in computational image analysis arising from very low EpiStain signals in some Epi^+^ cells of EMT^+^ tumors, we applied visual scoring for these markers. We used hematoxylin and eosin (H&E) and PAX8 staining on consecutive sections to confirm the presence of tumor epithelial cells.

Interestingly, among the tumors classified as CD45^high^, those with a low EpiStain phenotype displayed FAP^+^ ccRCC cells at the tumor border more frequently (72%) compared to tumors with medium (39%) or high (32%) EpiStain phenotypes (Pearson chi-square test, p < 0.001; **Table 1 and Fig. 3H–3K**). This association was not as strong when FAP positivity was analyzed in the tumor center cores (Pearson chi-square test, p = 0.052; **Supplementary Table S3**). Automated digital quantification of FAP fluorescence intensity within Epi^+^ cells across EpiStain categories confirmed the visually observed differences at the tumor border (**Supplementary Fig. S4A**). To validate the specificity of the FAP antibody, we performed immunohistochemistry (IHC) using a CRISPR-Cas9 FAP knock-out model, which demonstrated complete loss of FAP staining in the knock-out cells (**Supplementary Fig. S4B**). Additionally, analysis of FAP expression at the RNA level in renal cell carcinoma cell lines indicated notable FAP mRNA expression in HCC89 carcinoma cells (**Supplementary Fig. S4C, S4D**). This additional validation strongly supports the specificity of the FAP antibody and confirms its expression in ccRCC tumor cells.

**Table 1.** Epithelial marker expression association with mesenchymal protein expression in tumor cells in ccRCC with high leukocyte infiltration (CD45^high^).

| Tumor border variable<sup>b</sup> | EpiStain intensity category<sup>a</sup> |  |  | p |
| --- | --- | --- | --- | --- |
|  | Low (n=45) | Med (n=45) | High (n=45) |  |
| **FAP (tumor)** |  |  |  | **<0.001** |
| Neg (n=71) | 13 (28 %) | 31 (61 %) | 23 (68 %) |  |
| Pos (n=58) | 34 (72 %) | 20 (39 %) | 11 (32 %) |  |
| na (n=3) |  |  |  |  |
| **SPARC (tumor)** |  |  |  | **0.001** |
| Neg (n=74) | 16 (36 %) | 36 (71 %) | 22 (67 %) |  |
| Pos (n=55) | 29 (64 %) | 15 (29 %) | 11 (33 %) |  |
| na (n=6) |  |  |  |  |
| **VIM (tumor)** |  |  |  | **0.018** |
| Neg (n=21) | 2 (4 %) | 13 (25 %) | 6 (18 %) |  |
| Pos (n=108) | 43 (96 %) | 38 (75 %) | 27 (82 %) |  |
| na (n=6) |  |  |  |  |
| **PD-L1 (tumor)** |  |  |  | **0.002** |
| Neg (n=81) | 20 (46 %) | 33 (65 %) | 28 (85 %) |  |
| Pos (n=46) | 23 (54 %) | 18 (35 %) | 5 (15 %) |  |
| na (n=8) |  |  |  |  |

<sup>a</sup>EpiStain intensity was measured in Epi<sup>+</sup> cells in tumor center cores (2 replicates) and CD45<sup>high</sup> patients (n = 135) were categorized into three equally sized groups based on the mean tumor center core intensities.
<sup>b</sup>Mesenchymal proteins were visually scored in tumor border TMA cores (2 replicates for each patient). Positive staining in tumor cells in either replicate core was evaluated as a positive score. p-value, Pearson Chi-square (Exact two-sided). See also Supplementary Table S3 for tumor center analysis.

To further investigate EMT charateristics in CD45^high^ tumors, other mesenchymal markers—such as secreted protein acidic and rich in cysteine (SPARC) and vimentin (VIM)—as well as PD-L1 were more commonly expressed in ccRCC cells of tumors with lower EpiStain levels in border regions (Pearson chi-square test, p = 0.001, p = 0.018, and p = 0.002, respectively; **Table 1**). Consistent with recent findings that mesenchymal gene induction occurs at invasive tumor borders in ccRCC,(26,28) we observed that FAP and VIM were both more frequently positive in ccRCC cells at the tumor border compared to tumor center areas (FAP: 49.6% vs. 25.2%; VIM: 83.7% vs. 69.4%).

Extending this analysis to the entire ccRCC cohort, we found that high tumor cell-specific FAP expression correlated with increased CD45^+^ cell densities (Pearson chi-square test, p < 0.001), decreased CD31^+^ cell densities (p < 0.001), and higher levels of tumor PD-L1 expression (p < 0.001) (**Supplementary Table S4**). Notably, other mesenchymal markers studied (PDGFRA, PDGFRB, αSMA, POSTN) did not show epithelial positivity (data not shown). Tumors with a CD45^high^EMT^+^ phenotype—marked by higher tumor cell FAP levels—were associated with increased PD-L1 expression in tumor cells (Pearson chi-square test, p < 0.001), a pattern not associated with SPARC or VIM (**Fig. 3L and Supplementary Table S5**).

Kaplan-Meier analysis revealed that higher FAP positivity in tumor cells correlates with shorter RFS in both the tumor border and center areas within the CD45^high^ group (**Fig. 3M-3N**), as well as in the overall patient population (**Supplementary Fig. S4E-S4F**). However, despite its correlation with FAP positivity, tumor PD-L1 expression was not associated with RFS when scored in a similar manner as FAP (**Fig. 3O-3P; Fig. S4G–S4H;** see Materials and Methods for scoring strategy). Neither tumor SPARC nor VIM showed any association with RFS in either the tumor border or tumor center areas (**Supplementary Fig. S5A-S5D**).

Interestingly, ccRCCs with stronger tumor cell FAP expression—but not PD-L1 expression—were significantly more likely to metastasize to the liver compared to other locations (chi-square test, p < 0.001; **Supplementary Fig. S6A-D**). This association remained significant even after adjusting for clinical covariates, with an odds ratio (OR) of 2.08 (p = 0.017) for tumors with high tumor cell FAP expression (**Supplementary Fig. S6E**).

In summary, our analysis highlights a strong association between the loss of epithelial differentiation markers and increased tumor cell FAP expression, especially at the ccRCC tumor border. This suggests that FAP marks an aggressive cancer cell EMT status distinct from other EMT markers, such as SPARC and VIM, and is superior to PD-L1 in risk stratifying localized ccRCC, with tumor FAP expression correlating strongly with liver metastasis.

### FAP⁺ cancer-associated fibroblasts are enriched at tumor edges in CD45^high^, EMT⁺ ccRCC

We next examined the role of cancer-associated fibroblasts (CAFs) located at the tumor edge, focusing on their association with the loss of epithelial markers in tumors with high CD45⁺ cell infiltration (CD45^high^ tumors). This exploration is key to understanding the connection between stromal cell subsets and tumor cell phenotypes within the context of EMT. Using the CAF panels for our analysis, we observed that out of eight single-marker-defined stromal cell subsets, only FAP⁺ stromal cells (FAP⁺Epi⁻) exhibited a statistically significant increase in abundance at the tumor border, correlating with a reduction in EpiStain intensities from high to medium and low levels in CD45^high^ tumors (Kruskal-Wallis test, p = 0.035; **Fig. 4A**). The abundance of PAX8⁺ epithelial cells did not vary significantly among cases with low, medium, and high EpiStain levels (**Fig. 4B**), suggesting that this epithelial marker is better retained during the EMT process than E-cadherin, cytokeratins, and CA9 (components of the EpiStain cocktail). In our examination of the entire ccRCC cohort, we observed a notable association between the abundance of FAP⁺ stromal cells at the tumor border and FAP⁺ tumor cells (Kruskal-Wallis test; p < 0.001; **Fig. 4C**), as well as a positive relationship with overall leukocyte counts (Kruskal-Wallis test; p < 0.001; **Fig. 4D**).

Building on these observations of stromal cells defined by single markers, we next explored the expression patterns of stromal cells identified by combinations of multiple markers (CAF subsets; see **Fig. 4E-4F** for classes and distributions). This multi-marker approach effectively identifies stromal or fibroblast cells, reducing the likelihood of misidentifying individual markers—for example, FAP in epithelial cells, PDGFRB in pericytes, and αSMA in smooth muscle cells. Interestingly, we found that the relative proportions of multi-marker defined CAFs were remarkably consistent between the tumor border and tumor center in all ccRCC cases (**Fig. 4E and 4F**). Importantly, our observations revealed a strong association between poor recurrence-free survival (RFS) and the presence of FAP⁺ stromal cells. This association was evident both in cases marked by FAP positivity alone and in those where FAP was combined with PDGFRB (CAF6) and αSMA (CAF7), particularly at the tumor border (Cox regression; p < 0.05; **Fig. 4G and 4H**). FAP⁺ stromal cells were also prognostically significant within tumor center cores (Cox regression; p < 0.05; **Fig. 4H**). Conversely, markers in CAF panel 2 (SPARC, PDGFRB, POSTN, VIM), individually or in combinations (CAF16 to CAF30) in the tumor border, showed no associations with RFS (**Fig. S7A and S7B**). Within the tumor center cores, we found that POSTN⁺ stromal cells (see **Fig. 4I** for example), alone or in combination with PDGFRB and SPARC (CAF29), were associated with improved RFS, while the POSTN⁻ CAF24 subset (VIM⁺SPARC⁺POSTN⁻PDGFRB⁻) correlated with less favorable RFS (**Fig. S7C and S7D**). Collectively, our findings reveal a pronounced enrichment of FAP⁺ stromal cells at the tumor border in CD45^high^, EMT⁺ tumors, establishing them as significant prognostic markers. Interestingly, this finding contrasts with the observations for POSTN⁺ cells, which are associated with favorable survival in the tumor center, highlighting how different fibroblast markers can have region-specific effects within the ccRCC tumor microenvironment.

### FAP^+^EMT^+^ status is associated with an immunosuppressive microenvironment

To further characterize the immunological landscape of the EMT phenotype in ccRCC, we analyzed the proportions of diverse lymphoid and myeloid cell subsets within CD45^high^ tumors, stratified by low, medium, and high EpiStain intensities. We observed no significant differences in T-cell densities (CD3^+^CD45^+^) or other lymphoid cells (T cells, NK cells, B cells) relative to the total CD45^+^ population across the EpiStain categories (**Fig. 5A**). However, within the myeloid cell population, CD11c^+^ cells (CD11c^+^CD45^+^ and CD68^-^ CD11c^+^CD45^+^), typically marking dendritic cells, showed an increase from low to high EpiStain cases in the CD45^high^ tumors (Kruskal-Wallis test, p < 0.001 and p = 0.009, respectively; **Fig. 5A**).

The ratio of M2-like macrophages (CD68^+^CD163^+^CD45^+^) to M1-like macrophages (CD68^+^CD11c^+^CD45^+^) was found to be lower in cases with high EpiStain levels (mean ratio 1.47) compared with low (ratio 4.70) or medium (ratio 4.33) EpiStain levels (Kruskal-Wallis test, p = 0.001; **Fig. 5B**), suggesting an enrichment of immunosuppressive macrophages in CD45^high^ tumors with loss of EpiStain markers. In analyzing T-cell subsets, we found an accumulation of terminally exhausted T effector cells (PD1^+^TIM3^+^CD8^+^CD3^+^) with decreasing EpiStain levels in tumor center (Kruskal-Wallis test, p = 0.008; **Fig. 5C**) and border cores (p = 0.008; **Supplementary Fig. S8A**), but not in benign cores (p = 0.550; **Supplementary Fig. S8B**). No differences were observed in the densities of granzyme B-positive (GrzB^+^CD8^+^CD3^+^) or PD-L1-positive (PD-L1^+^CD8^+^CD3^+^) T-cell subsets within the EpiStain low, medium, and high tumor categories (Kruskal-Wallis test, p = 0.463 and p = 0.250, respectively; **Fig. S8C and S8D**). T-regulatory cells (Tregs, FOXP3^+^CD8^-^CD3^+^) showed a trend toward higher densities with lower EpiStain levels in tumor center cores (Kruskal-Wallis test, p = 0.091; **Fig. S8E**). Importantly, the immunosuppressive exhausted T-cell phenotype (PD1^+^TIM3^+^CD8^+^CD3^+^) was also enriched in patients with stronger tumor cell FAP both in tumor center and border areas in the CD45^high^ group and in the overall patient population (Kruskal-Wallis test, p < 0.001; **Fig. 5D and S8G**). Additionally, tumor border Tregs (FOXP3^+^CD8^-^CD3^+^) and M2-like macrophages (CD68^+^CD163^+^CD45^+^) were more prominent in cases with higher tumor FAP levels both in the CD45^high^ group and in the overall patient population (**Fig. 5D and 5E and S8H**). This comprehensive analysis underscores the increased presence of immunosuppressive immune cell phenotypes with higher tumor FAP levels and loss of epithelial markers in ccRCC patients with poor RFS.

### FAP predicts poor survival in early-stage and sunitinib-treated ccRCC patients

In the treatment landscape of localized clear cell renal cell carcinoma (ccRCC), there is a notable divergence in therapeutic strategies between early-stage (pT1–2) and late-stage (pT3–4) patients. Current molecular and cellular profiling studies primarily focus on refining treatment stratification for late-stage patients, especially for tyrosine kinase inhibitors (TKIs) and immune checkpoint blockade (ICB) therapies. For survival analysis, we focused on FAP and EMT-associated phenotypes in early-stage ccRCC patients (pT1–2; n = 303). Cox regression analysis with continuous variables revealed that higher counts of stromal FAP⁺ cells, stronger tumor FAP expression, and increased CD45⁺ cell density are significantly associated with poorer recurrence-free survival (RFS) in early-stage localized ccRCC (**Fig. 6A**). Conversely, a higher density of CD31⁺ cells correlated with improved RFS in these patients (**Fig. 6A**).

**Figure 6.**
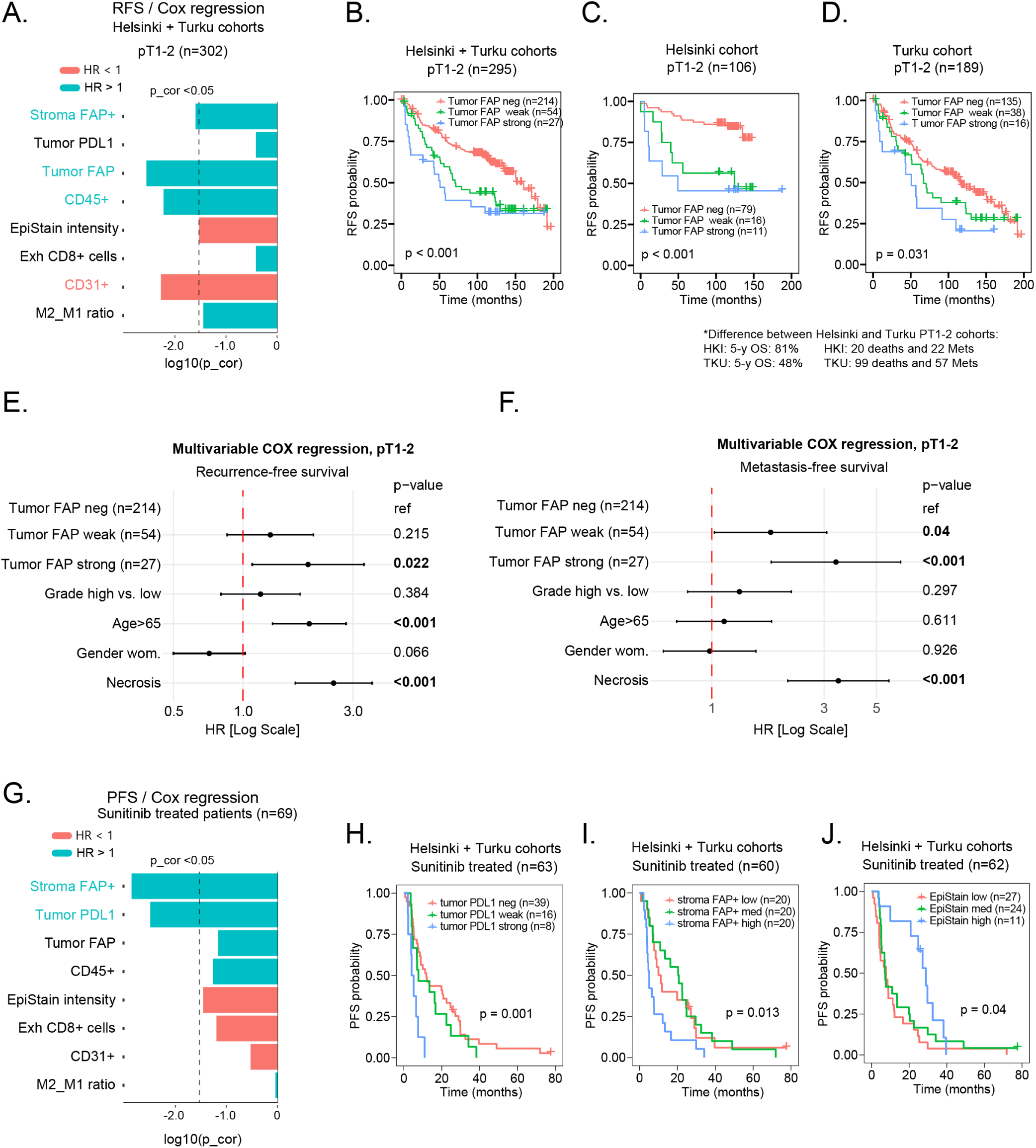
FAP predicts survival in early-stage and sunitinib-treated ccRCC patients. **(A)** Univariate Cox regression analysis for early-stage ccRCC patients, using continuous variables. HR, Hazard ratio; p_cor, Bonferroni corrected p-value. Exh CD8^+^, Exhausted T cells (PD1^+^TIM3^+^CD3^+^CD8^+^/CD3^+^CD8^+^). **(B-D)** Kaplan-Meier survival analysis based on tumor-specific FAP expression (3-tier visual scoring) in early-stage ccRCC. Analysis for combined cohorts (B), Helsinki cohort (C), and Turku cohort (D). FAP positivity determined from maximum expression in tumor center or border. p-values from log-rank test. **(E, F)** Multivariable Cox regression for recurrence-free survival (RFS) (E) and metastasis-free survival (MFS) (F) in early-stage patients, considering multiple variables. (G) Univariate Cox regression for sunitinib-treated ccRCC patients (n = 69), with progression-free survival (PFS) as the outcome. HR, Hazard ratio; p_cor, fdr-corrected p-value. **(H-J)** Kaplan-Meier curves evaluating tumor-specific PDL1 expression (H), stromal FAP^+^ cell density (I), and EpiStain intensity (J) in patients treated with sunitinib after detection of metastases. PFS, progression-free survival. p-values from log-rank test.

Kaplan-Meier analysis, focusing on categorized tumor FAP expression levels (negative, low, strong) in the combined cohort, further validated the strong association of tumor FAP expression with poor prognosis among pT1–2 patients (log-rank test, p < 0.001; **Fig. 6B**). This association was consistently observed in both the Helsinki (p < 0.001; **Fig. 6C**) and Turku sub-cohorts (p = 0.031; **Fig. 6D**). In pT1–2 patients, strong tumor FAP expression emerged as an independent predictor of outcome, even after adjusting for grade, age, gender, and necrosis (Cox regression, p = 0.022; **Fig. 6E**). This prognostic impact was more pronounced when using metastasis-free survival (MFS) as the endpoint rather than recurrence (Cox regression, p < 0.001; **Fig. 6F**).

Notably, patient age lost its significance as a prognostic factor with MFS, suggesting that while age may influence mortality, tumor FAP is a more potent determinant of metastasis risk.

For comparative purposes, we then assessed the same primary tumor phenotypes for their associations with progression-free survival (PFS) in patients treated with first-line sunitinib after developing metastases (n = 69). In this group, increases in stromal FAP⁺ cell counts or PD-L1⁺ tumor cells were linked to worse PFS (**Fig. 6G– 6I**). Conversely, EpiStain intensity, when analyzed as a continuous or categorized variable, showed a trend toward improved PFS (**Fig. 6G and 6J**).

Considering the known association of sarcomatoid RCC with increased tumor PD-L1 expression, enhanced immune infiltration, and an EMT phenotype (15,40). we explored whether tumor and stromal FAP similarly mark ccRCC cases with sarcomatoid features. In our cohort, associations were observed between sarcomatoid morphology and tumor PD-L1, tumor FAP, stromal FAP, but not with CD45⁺ cell infiltration or EpiStain intensity (**Supplementary Table S6**). Notably, among ccRCCs with high CD45⁺ cell infiltration, sarcomatoid morphology correlated with PD-L1 expression but not with tumor/stroma FAP positivity (**Supplementary Table S7**). This suggests that high CD45⁺ infiltration alongside elevated tumor and stromal FAP expression identifies a unique ccRCC subtype, distinct from established features of sarcomatoid transformation.

## DISCUSSION

### Context and objectives of the study

In this study, we utilized multi-region single-cell multiplexed immunofluorescence to explore the phenotypic heterogeneity of clear cell renal cell carcinoma (ccRCC). Our primary aim was to investigate the tumor microenvironment (TME) and tumor cell phenotypes in ccRCC cases with high immune infiltration (CD45^high^). This focus arises from the need to better understand the role of immune cells and their interactions with other TME components and tumor cell features in ccRCC, as these interactions are critical in driving disease progression and potentially influencing the efficacy of systemic therapies such as tyrosine kinase inhibitors (TKIs) and immune checkpoint blockade (ICB). We specifically concentrated on localized ccRCC, where the nature and implications of immune infiltration have been less explored compared to metastatic ccRCC.

We identified a distinct ccRCC subtype characterized by high immune infiltration, immunosuppression, and an epithelial-to-mesenchymal transition (EMT) phenotype in two independent cohorts of localized tumors. Notably, this subtype exhibits dual expression of fibroblast activation protein (FAP) in both tumor and stromal cells and is associated with poor recurrence-free survival (RFS) after radical or partial nephrectomy, as well as reduced response to first-line sunitinib treatment after the development of metastases.

### The prognostic implications of immune infiltration and angiogenesis in ccRCC

Our investigation of localized ccRCC demonstrated that increased CD45^+^ cell density—reflecting robust immune infiltration in various tumor areas, including the tumor center, edge, and adjacent benign regions—was associated with worse recurrence-free survival (RFS). Conversely, elevated densities of CD31^+^ endothelial cells, indicative of angiogenesis, were linked to improved RFS.

Our single-cell spatial analysis significantly expands on recent gene expression signature studies linking angiogenesis to better survival in metastatic ccRCC (12–15,41) and localized ccRCC (17). Additionally, our findings support previous observations that immune infiltration—especially involving exhausted T cells and myeloid cell populations—is associated with poorer survival (13,17,20,22). These cells can create an immunosuppressive milieu that facilitates tumor evasion from immune surveillance and enhances metastatic potential (42).

### The role of FAP and EMT in tumor progression and prognosis

In our study, we classified ccRCC cells according to their epithelial marker characteristics using an EpiStain antibody cocktail comprising antibodies against CA9, E-cadherin, and cytokeratins. This cocktail was designed to detect a wide variety of epithelial cells in the tumor center, edge, and benign areas. E-cadherin and cytokeratins are epithelial differentiation proteins often lost during epithelial-to-mesenchymal transition (EMT) (43). CA9, which has not been well characterized in the context of EMT in ccRCC, can also exhibit highly heterogeneous expression in ccRCC cells, as shown by Davidson et al. (28) and our re-analysis of the scRNA-seq dataset from Li et al. (26).

We observed significant variation in EpiStain intensity within the tumor cells of localized ccRCC. In CD45^high^ tumors, lower EpiStain levels in ccRCC cells were associated with poorer RFS, decreased CD31^+^ endothelial cell density, and increased mesenchymal marker expression (VIM, SPARC, FAP) within tumor cells at the tumor border. This is indicative of EMT or partial/hybrid EMT, as other mesenchymal markers were negative and the PAX8 epithelial marker did not show significantly reduced levels in these Epi^+^ cells.

Tumors characterized by EpiStain loss and tumor FAP expression often exhibited PD-L1 positivity, linking EMT to immune modulation. Tumor-specific FAP expression was associated with worse RFS in both the overall ccRCC cohort and the CD45^high^ subgroup. Gene signature studies (15,17) and single-cell profiling (26,28) have emphasized the role of EMT in ccRCC progression. Our findings extend this knowledge by showing FAP— typically a cancer-associated fibroblast (CAF) marker—expressed in tumor cells of the aggressive CD45^high^ ccRCC subtype. Similar associations between tumor-specific FAP and EMT or pro-invasiveness have been reported in melanoma,(44) hepatocellular carcinoma (45), and ovarian cancer (46).

### The significance of tumor cell phenotypes and their interactions with cancer-associated fibroblasts

In our study of localized ccRCC, we identified a CD45^high^EMT^+^ phenotype associated with increased FAP^+^ cancer-associated fibroblasts (CAFs) at tumor borders. The density of these FAP^+^ CAFs correlated with both tumor FAP levels and immune cell infiltration in the total patient population. Notably, the presence of FAP^+^ CAFs predicted shorter RFS in our localized ccRCC cohort, both when measured in tumor center and edge regions. This emphasizes the critical role of tumor-stromal interactions in ccRCC progression.

We also explored multi-marker-defined CAFs, extending insights from our recent mIF profiling study in non-small cell lung cancer (NSCLC) (37). In NSCLC, we identified a particularly aggressive subset of FAP^+^ CAFs (CAF7: αSMA^+^/FAP^+^/PDGFRB^+^/PDGFRA^-^), associated with *TP53* mutations, tumor PD-L1 positivity, and infiltration of M2-like tumor-associated macrophages (TAMs) (37). In ccRCC, this CAF7 subset was prognostically significant in tumor edge regions, aligning with findings from Davidson et al. (28), who reported a similar myCAF subset (αSMA^+^, FAP^+^, FN1^+^) enriched at EMT^+^ ccRCC tumor borders. They further demonstrated through analysis of The Cancer Genome Atlas (TCGA) ccRCC RNA-seq data (KIRC) that these myCAFs were associated with the presence of exhausted CD8^+^ T cells and TAMs. This aligns with our observations, highlighting the crucial role of CAFs in shaping the immune landscape of ccRCC.

Furthermore, Motzer et al. (15) classified ccRCCs into seven RNA-seq-based molecular subtypes and identified a “stroma/proliferating” gene expression signature marked by CAF markers such as FAP, FN1, and POSTN (periostin), coupled with low angiogenesis and high EMT, which correlated with poor prognosis. In contrast, Davidson et al. (28) demonstrated that POSTN was highly co-expressed with MHC class II transcripts (HLA-DRs), defining antigen-presenting CAFs (apCAFs) associated with better overall survival. Consistent with their study, our single-cell spatial analysis found that a higher number of POSTN^+^ stromal cells—and the specific multi-marker-defined subset POSTN^+^PDGFRB^−^SPARC^−^VIM^−^ (CAF17) in tumor center areas, but not in tumor borders—were associated with improved RFS. These studies emphasize that CAFs with distinct molecular profiles can have divergent roles in tumor progression, underlining the need for detailed spatial profiling to understand these complex interactions.

### Immune cell subsets enriched in CD45^high^EMT^+^ ccRCC

In analyzing CD45^high^ ccRCCs, we observed increased CD11c^+^ dendritic cell densities in tumors with higher EpiStain levels. In contrast, exhausted T cells (CD3^+^CD8^+^PD-1^+^TIM-3^+^) and M2-like tumor-associated macrophages (TAMs; CD163^high^/CD11c^low^) were more prevalent in tumors exhibiting loss of EpiStain and EMT^+^ phenotype. The accumulation of terminally exhausted T effector cells, regulatory T cells (Tregs), and M2-like macrophages in tumors with higher tumor/stromal FAP expression further underscores the tight associations among immune suppression, EMT, and tumor progression in ccRCC.

These findings align with studies showing enriched exhausted T cells and M2-like TAMs in advanced ccRCC stages (19) and the presence of specific IL1B-positive macrophages in proximity to EMT^high^ ccRCC cells at invasive tumor borders (26). Furthermore, a large-scale RNA-seq study from the PROTECT trial indicated that EMT and myeloid, rather than lymphoid, inflammation is associated with faster metastatic recurrence in high-risk, localized ccRCC (17).

### EMT-associated phenotypes in ccRCC and their prognostic implications with potential for early intervention

Our findings significantly enhance the understanding of highly immune-infiltrated and EMT-associated phenotypes in localized ccRCC. Notably, we highlight a specific EMT phenotype characterized by tumor cell FAP expression. This high FAP expression correlates with poorer outcomes, even in a subgroup of CD45^high^ or early-stage (pT1–T2) patients, suggesting its potential as a prognostic marker and therapeutic target.

Furthermore, we found that high tumor FAP expression is specifically associated with an increased incidence of liver metastases in ccRCC patients, implying that tumor cell FAP may play a role in promoting metastatic spread to the liver. Our study also demonstrates that higher FAP expression in stromal cells is associated with a reduced benefit from first-line sunitinib treatment after onset of metastasis. This suggests that FAP plays a significant role not only in the primary tumor microenvironment but also potentially in metastatic settings, especially in the liver. Interestingly, a recent report on colorectal cancer liver metastasis suggests that stromal FAP in liver metastasis models promotes resistance to anti-angiogenic therapy by inducing tumor cell EMT and recruiting myeloid cells into the microenvironment (47). These findings indicate that similar biological mechanisms—such as EMT induction and myeloid cell recruitment—underlie both primary tumor progression and metastatic colonization in the liver, with FAP serving as a potential mediator. This suggests that FAP-linked pathways governing EMT and myeloid cell recruitment contribute to resistance against angiogenic therapies like sunitinib in both primary tumors and metastatic sites such as the liver. Understanding these shared mechanisms opens avenues for developing targeted therapies that inhibit FAP, potentially overcoming treatment resistance and preventing liver metastasis in ccRCC patients. Such strategies could be integrated with existing angiogenic therapies to enhance their efficacy and improve patient outcomes.

Our results suggest that in the overall ccRCC cohort, FAP⁺EMT⁺ ccRCCs share characteristics with sarcomatoid RCC, which is known for having more frequent tumor PD-L1 expression, low angiogenesis, high immune infiltration, a prominent stroma/proliferating signature, EMT features, and poorer survival (15,40). However, when focusing on the CD45^high^ subgroup, we found that the association between FAP⁺EMT⁺ phenotype and sarcomatoid features was not observed. Specifically, among the CD45^high^ tumors, 11 out of 131 tumors (8.4%) exhibited sarcomatoid differentiation. Within this subgroup, only tumor PD-L1 expression was significantly associated with sarcomatoid morphology, while the aggressive FAP⁺ phenotype did not show such an association. This indicates that, within the tumors with high leukocyte infiltration (CD45^high^), the aggressive FAP^+^ phenotype represents a distinct subtype of ccRCC.

Sarcomatoid ccRCC is often enriched with mutations in *BAP1*, *PTEN*, *TP53*, and *CDKN2A*/*B*, while typically exhibiting fewer *PBRM1* mutations (15,17,40). Investigating the relationship between FAP-expressing tumor cells, CAFs, and these genetic mutations could be highly informative, especially considering our previous observation of a correlation between the FAP^+^ CAF7 subset and *TP53* mutations (37). Understanding these connections in tumors with both high and low leukocyte infiltration may shed light on the intricate interplay between genetic alterations and the tumor microenvironment (TME) in ccRCC.

Dual expression of FAP in tumor cells and CAFs, associated with increased immune infiltration, may potentially serve as a biomarker for predicting not only ccRCC response to angiogenic therapies but also to ICB. This is particularly relevant considering the enhanced responsiveness of sarcomatoid ccRCC subtypes to ICB treatment (5,15) and their lower response to TKI treatment (5). However, a hypothesis that EMT-associated dual positivity of FAP in tumor and stromal cells could mark ccRCCs with a better response to ICB is challenged by findings from Davidson et al. (28), who analyzed bulk RNA-seq data from the BIONIKK clinical trial involving advanced ccRCC patients treated with nivolumab. Their study suggested that an mRNA signature indicating a high association of EMT^high^ ccRCC cells (ccRCC.mes) with myCAFs (FAP^+^ subset) was associated with resistance to ICB therapy. However, our study strongly suggests that the risk and treatment prediction associations with the TME and cancer cell states should be correlated with leukocyte infiltration rates in ccRCC—an aspect not addressed in the Davidson et al. (28) study. Therefore, integrating these observations of leukocyte infiltration, stromal components, and cancer cell states with genomic profiling in future studies will enable us to better stratify patients for tailored treatment strategies.

### Potential FAP-targeted therapeutic implications and future directions

FAP-targeted imaging and therapeutic strategies present exciting opportunities in ccRCC, particularly due to the dual expression of FAP in both cancer and stromal cells. FAPI-PET imaging, a novel approach that targets FAP, offers the potential to non-invasively identify aggressive ccRCC tumors by visualizing FAP expression (48). Our findings suggest that FAPI-PET imaging could be explored for early detection of high-risk ccRCC, aiming to improve patient stratification and inform treatment decisions—for example, identifying patients in need of neoadjuvant or adjuvant therapy.

Additionally, FAP-targeted radioligand therapy and bispecific antibodies could be used to eliminate FAP-expressing cells, offering novel ways to treat these tumors (49). By simultaneously targeting the FAP-positive tumor microenvironment and stimulating the immune system (50,51), these therapies could be particularly effective in aggressive ccRCC subtypes characterized by high immune infiltration and dual FAP expression.

### Limitations and conclusions

The limitations of this study include its retrospective design. While the patients had undergone radical or partial nephrectomy before pembrolizumab was introduced as an adjuvant treatment, we were unable to analyze the impact of ccRCC cancer cell states and TME characteristics on the outcomes of adjuvant immunotherapy. This will be the focus of our future studies.

In conclusion, our study elucidates the complex interplay between immune cell infiltration, FAP expression, and EMT in the progression of localized ccRCC. A pivotal finding was the dual presence of FAP in both tumor and stromal cells, especially within the context of EMT, linking high FAP expression to poorer prognosis. Moreover, the correlation of elevated tumor PD-L1 expression and increased stromal FAP^+^ CAFs with the EMT^+^ primary tumor phenotype underscores their combined negative impact on the efficacy of first-line sunitinib treatment after the development of metastases.

Our extensive analysis highlights the significance of FAP in the progression of localized ccRCC, including in early-stage (pT1–T2) cases with high immune infiltration. Future research should build on these insights to guide biomarker selection for enhancing therapeutic decision-making and improving neoadjuvant and adjuvant treatments for patients with localized ccRCC.

## AUTHORS’ CONTRIBUTIONS

T.P. conceived and supervised the study, designed the experiments and the antibody panels, generated the mIF dataset, analyzed mIF data together with L.L., and wrote the manuscript with inputs from L.L., K.E.M., L.P., T.M., P.M.J., O.B., P.V., S.V., O.K. A.H. and K.V. helped with the antibody panel design, antibody testing, and performed the staining and imaging. T.M., L.L., P.V., K.E.M., P.M.J., H.N., P.J., E.K. provided the clinical and pathological data and gave clinical input. L.C. performed the scRNA-seq analysis.

## ACKOWLEDGEMENTS

We thank the FIMM Digital Microscopy and Molecular Pathology Unit supported by University of Helsinki and Biocenter Finland. T.P. thanks the financial support from the Sigrid Juselius Foundation (Decisions 230149, 240158), Cancer Foundation Finland (Decision date, 30/11/2022), and Research Council of Finland (Decision no. 363152). For this study, T.M. has been funded by Research Council of Finland, Hospital District of Helsinki and Uusimaa, Cancer Foundation Finland. The graphical abstract was created with BioRender (biorender.com).

L.L acknowledges Hospital District of Helsinki and Uusimaa, Finnish Urological Association (Urologiyhdistys ry), Maud Kuistila grant and Finnish Urological research foundation.

