## Supplemental Data for "Fibroblast Activation Protein Defines an Aggressive, Immunosuppressive EMT-associated Tumor Subtype in Highly Inflamed Localized Clear Cell Renal Cell Carcinoma"


**Supplementary Table S1:** Clinicopathological characteristics of patients.

**Supplementary Table S2**: Antibody panel details.

**Supplementary Table S3**: EpiStain intensity association with mesenchymal protein expression in tumor cells (center) in ccRCC with high leukocyte infiltration ($CD45^{high}$).

**Supplementary Table S4**: Tumor FAP association with EpiStain levels, leukocyte counts, tumor PD-L1, and endothelial cell counts.

**Supplementary Table S5**: Tumor PD-L1 association with tumor FAP and EpiStain levels in $CD45^{high}$ ccRCC.

**Supplementary Table S6**: Tumor PD-L1 and FAP association with sarcomatoid ccRCC in full cohort.

**Supplementary Table S7**: Tumor PD-L1 and FAP association with sarcomatoid ccRCC in patients with high tumoral leukocyte infiltration ($CD45^{high}$).

**Supplementary Fig. S1**: Kaplan-Meier plots of RFS for patients stratified by tertiles of $CD45^{+}$ cell counts.

**Supplementary Fig. S2**: Single-cell scatter plot of VIM and SPARC expression in RCC tumor cells with CA9 expression annotation.

**Supplementary Fig. S3**: Correlation plots of EpiStain intensity in $Epi^{+}$ cells between tumor center, border, and adjacent benign TMA cores.

**Supplementary Fig. S4**: Association of tumor cell FAP and PD-L1 expression with recurrence-free survival (RFS) in tumor center cores of localized ccRCC.

**Supplementary Fig. S5**: Tumor cell SPARC and VIM association with RFS in localized $CD45^{high}$ ccRCC.

**Supplementary Fig. S6**: Comparative analysis of FAP and PD-L1 expression across different metastasis locations in ccRCC tumors and their relationship with liver metastasis.

**Supplementary Fig. S7**: Distribution and survival associations of CAFs in localized ccRCC using markers from CAF panel 2.

**Supplementary Fig. S8.** Distribution of immune cell subsets across EpiStain and tumor FAP categories in localized ccRCC.

**Table S1**: Clinicopathological characteristics of patients.

| | Cohorts | |
|---|---|---|
| **Variable** | **Helsinki** | **Turku** |
| No. of patients | 196 | 239 |
| Median follow-up time[a], years (IQR) | 8.69 (2.08–10.23) | 6.01 (2.09–9.86) |
| 5-year RFS rate | 62% | 62% |
| 5-year OS rate | 81% | 73% |
| Age at diagnosis, mean | 64 | 67 |
| Sex | | |
| Male | 99 (51%) | 142 (59%) |
| Female | 97 (49%) | 97 (41%) |
| pTNM | | |
| pT1a-b | 87 (44%) | 136 (57%) |
| pT2a-b | 26 (13%) | 53 (22%) |
| pT3-T4 | 83 (42%) | 50 (21%) |
| Grade[b] | | |
| 1 | 14 (7%) | 14 (6%) |
| 2 | 105 (54%) | 118 (49%) |
| 3 | 67 (34%) | 86 (36%) |
| 4 | 10 (5%) | 15 (6%) |
| na | | 6 (3%) |
| Necrosis | | |
| No | 132 (67%) | 135 (56%) |
| Yes | 58 (30%) | 103 (43%) |
| na | 6 (3%) | 1 (0%) |
| Sarcomatoid | | |
| No | 163 (83%) | 227 (95%) |
| Yes | 10 (5%) | 12 (5%) |
| na | 23 (12%) | |

Abbreviations: IQR, inter-quartile range; RFS, recurrence-free survival; OS, overall survival; na, not available

[a]Follow-up times and IQR were based on observed times to recurrence or last follow-up without adjustment for censored data.

[b]Grade is either Fuhrman or ISUP, depending on the period of tumor resection.

**Table S2:** Antibody panel details.

| | **TME panel** | |
|---|---|---|
| **First round staining** | **Antibody; used dilution** | **Product information** |
| TSA-488 | M-anti-D2-40; 1:100 | Dako; M3619 |
| TSA-555 | R-anti-PDGFRB; 1:100 | CST; 3169 |
| Alexa-647 | M-anti-CD56; 1:100 | Dako; M7304 |
| Alexa-750 | R-anti-CD11b; 1:100 | BioSB; 6441 |
| **Second round staining** | | |
| Alexa-647 | M-anti-CD20; 1:100 | Thermo; MS-340 |
| Alexa-750 | R-anti-CD3; 1:150 | Thermo; MA5-14482 |
| **Third round staining** | | |
| Alexa-647 | R-anti-CD31; 1:200 | Abcam; ab28364 |
| Alexa-750 | M-anti-CD45; 1:100 | Dako; M0701 |
| **Fourth round staining** | | |
| Alexa-647 | **EpiStain cocktail:**<br>R-anti-CAIX (CA9); 1:200<br>R-anti-E-Cadherin; 1:200<br>R-anti-panCK; 1:200 | <br>NovusBio; NB100-417<br>CST; 3195<br>Abcam; 9377 |
| Alexa-750 | M-anti-HLA-DR 1:5000 | Abcam; ab20181 |

| | **Tcell panel** | |
|---|---|---|
| **First round staining** | **Antibody; used dilution** | **Product information** |
| TSA-488 | M-anti-PD1; 1:100 | LSBio; B12784 |
| TSA-555 | R-anti-CD3; 1:750 | Thermo; MA5-14482 |
| Alexa-647 | R-anti-PDL1; 1:100 | CST; 13684 |
| Alexa-750 | M-anti-CD8; 1:300 | Dako; M7103 |
| **Second round staining** | | |
| Alexa-647 | R-anti-Tim-3; 1:100 | CST; 45208 |
| Alexa-750 | M-anti-FoxP3; 1:100 | Abcam; ab20034 |
| **Third round staining** | | |
| Alexa-647 | R-anti-GranzymeB; 1:200 | Abcam; ab4059 |
| Alexa-750 | M-anti-Ki67; 1:100 | Dako; M7240 |
| **Fourth round staining** | | |

| | | |
|---|---|---|
| Alexa-750 | **R-anti-PanEpi cocktail**<br>R-anti-E-Cadherin; 1:200<br>R-anti-panCK; 1:200 | <br>CST; 3195<br>Abcam; 9377 |

| | **CAF panel 1** | |
|---|---|---|
| **First round staining** | **Antibody; used dilution** | **Product information** |
| TSA-488 | R-anti-PDGRFB; 1:100 | CST; 3169 |
| TSA-555 | R-anti-PDGRFA; 1:100 | CST; 5249 |
| Alexa-647 | M-anti-aSMA; 1:200 | DAKO; M0851 |
| TSAbio-Str750 | R-anti-FAP; 1:2000 | Abcam; ab207178 |
| **Second round staining** | | |
| Alexa-647 | **R-anti-PanEpi cocktail**<br>R-anti-panCK; 1:200<br>R-anti-E-Cadherin; 1:200 | <br>Abcam; ab9377<br>CST; 3195 |

| | **CAF panel 2** | |
|---|---|---|
| **First round staining** | **Antibody; used dilution** | **Product information** |
| TSA-488 | G-anti-SPARC; 1:500 | R&D; AF941 |
| TSA-555 | R-anti-PDGRFB; 1:100 | CST; 3169 |
| Alexa-647 | R- anti -Periostin; 1:500 | Abcam; ab215199 |
| Alexa-750 | M- anti -Vimentin; 1:2000 | Dako; M0725 |
| **Second round staining** | | |
| - | - | - |
| Alexa-750 | **M-anti-PanEpi cocktail:**<br>M-anti-panCK; 1:200<br>M-anti-panCK; 1:200<br>M-anti-E-Cadherin; 1:200 | <br>Abcam; ab7753<br>Invitrogen; MA5-13156<br>BD; 610182 |

| | **Macrophage panel** | |
|---|---|---|
| **First round staining** | **Antibody; used dilution** | **Product information** |
| TSA 488 | R-anti-CD11c; 1:2000 | Abcam; ab52632 |
| TSA 555 | M-anti-CD206; 1:500 | Protein tech; 60143-1-Ig |
| AF647 | R-anti-CD16, 1:100 | CellMarque; 116R-14 |
| AF750 | M-anti-CD68; 1:100 | Cell Marque; 168M-94 |

| Second round staining | | |
|---|---|---|
| Alexa-647 | M-anti-CD45; 1:100 | Dako; M0701 |
| Alexa-750 | R-anti-CD163; 1:200 | Abcam; ab188571 |
| **Third round staining** | | |
| Alexa-647 | M-anti-HLADR; 1:5000 | Abcam; ab20181 |
| Alexa-750 | **R-anti-PanEpi cocktail**<br>R-anti-E-Cadherin; 1:200<br>R-anti-panCK; 1:200 | <br>CST; 3195<br>Abcam; 9377 |

**Table S3.** EpiStain intensity association with mesenchymal protein expression in tumor cells (center) in ccRCC with high leukocyte infiltration (CD45$^{high}$).

| | **EpiStain intensity category[a]** | | | |
|---|---|---|---|---|
| **Tumor center variable[b]** | **Low (n=45)** | **Med (n=45)** | **High (n=45)** | **p** |
| **FAP (tumor)** | | | | 0.052 |
| Neg (n=101) | 33 (69%) | 41 (79%) | 27 (77%) | |
| Pos (n=34) | 15 (31%) | 11 (21%) | 8 (23%) | |
| **SPARC (tumor)** | | | | **0.029** |
| Neg (n=80) | 21 (45%) | 34 (65%) | 25 (71%) | |
| Pos (n=54) | 26 (55%) | 18 (35%) | 10 (29%) | |
| na (n=1) | | | | |
| **VIM (tumor)** | | | | **0.018** |
| Neg (n=41) | 9 (19%) | 23 (44%) | 9 (26%) | |
| Pos (n=93) | 38 (79%) | 29 (56%) | 26 (74%) | |
| na (n=1) | | | | |
| **PD-L1 (tumor)** | | | | **<0.001** |
| Neg (n=101) | 21 (44 %) | 36 (69 %) | 31 (89 %) | |
| Pos (n=34) | 27 (56 %) | 16 (31 %) | 4 (11 %) | |

[a]EpiStain intensity was measured in Epi$^{+}$ cells in tumor center cores (2 replicates) and CD45$^{high}$ patients (n = 135) were categorized into three equally sized groups based on the mean EpiStain intensity in Epi$^{+}$ cells.

[b]Mesenchymal proteins were visually scored in tumor center TMA cores (2 replicates for each patient). Positive staining in tumor cells in either replicate core was evaluated as a positive score.

p-value, Pearson Chi-square (Exact two-sided).

**Table S4**: Tumor FAP association with EpiStain levels, leukocyte counts, tumor PD-L1, and endothelial cell counts.

| | Tumor FAP max expression[a] | | | |
|---|---|---|---|---|
| **Variable** | **Neg (n=282)** | **Weak (n=89)** | **Strong (n=47)** | **p** |
| **EpiStain (center)** | | | | **<0.001** |
| Low (n=135) | 73 (27%) | 35 (40%) | 27 (59%) | |
| Med (n=135) | 98 (36%) | 27 (31) | 11 (24%) | |
| High (n=135) | 102 (37%) | 25 (29%) | 8 (17%) | |
| na (n=13) | | | | |
| **PD-L1 (tumor max)[b]** | | | | **<0.001** |
| Neg (n=297) | 230 (82%) | 45 (51%) | 20 (43%) | |
| Weak (n=98) | 48 (17%) | 36 (40%) | 14 (39%) | |
| Strong (n=25) | 4 (1%) | 8 (9%) | 13 (28%) | |
| **CD45 count[c]** | | | | **<0.001** |
| Low (n=136) | 110 (40%) | 22 (25%) | 4 (9%) | |
| Medium (n=134) | 97 (36%) | 22 (25%) | 15 (32%) | |
| High (n=135) | 65 (24%) | 43 (50%) | 27 (59%) | |
| na (n=13) | | | | |
| **CD31 count** | | | | **<0.001** |
| Low (n=137) | 69 (25%) | 38 (44%) | 30 (65%) | |
| Medium (n=135) | 91 (34%) | 33 (38%) | 11 (24%) | |
| High (n=133) | 112 (41%) | 16 (18%) | 5 (11%) | |
| na (n=13) | | | | |

[a]Tumor FAP expression score as a maximum (max) score. FAP expression was scored as negative (score = 0), weak (score = 1), strong (score = 2) in all the tumor center/border TMA cores (2 x center; 2 x border for each patient tumor) and the highest score was selected as the patient-wise score in this analysis. n(tumor FAP core patients) = 418.

[b]PD-L1 tumor max scores were scored as for tumor FAP scores.

[c]CD45 and CD31 cell counts as fractions of Epi$^-$ cells analyzed in tumor center cores (mean of 2 replicates). The categorization into low, med, and high counts is based on the 3-tier equal categorization derived from the Helsinki and Turku ccRCC cohorts' categorization separately before cohort merging. This explains the slightly different n values for each category.

p-value, Pearson Chi-square (Exact two-sided).

**Table S5**: Tumor PD-L1 association with tumor FAP and EpiStain levels in CD45$^{high}$ ccRCC.

| | **Tumor PD-L1 max expression[a]** | | | |
|---|---|---|---|---|
| **Variable** | **Neg (n=73)** | **Weak (n=47)** | **Strong (n=15)** | **P** |
| **EpiStain (center)[b]** | | | | **<0.001** |
| Low (n=45) | 17 (23.3%) | 19 (40.4%) | 9 (60.0%) | |
| Med (n=45) | 20 (27.4%) | 19 (40.4) | 6 (40.0%) | |
| High (n=45) | 36 (49.3%) | 9 (19.1%) | 0 (0.0%) | |
| **FAP (tumor any)[c]** | | | | **<0.001** |
| Neg (n=70) | 43 (58.9%) | 20 (42.6%) | 2 (13.3%) | |
| Pos (n=64) | 29 (39.7%) | 27 (57.4) | 13 (86.7%) | |
| na (n=1) | 1 (1.4%) | 0 (0.0%) | 0 (0.0%) | |
| **SPARC (tumor any)** | | | | 0.463 |
| Neg (n=71) | 40 (54.8%) | 20 (42.6%) | 7 (46.7%) | |
| Pos (n=58) | 33 (45.2%) | 27 (57.4) | 8 (53.3%) | |
| **VIM (tumor any)** | | | | 0.254 |
| Neg (n=16) | 11 (15.1%) | 5 (10.6%) | 0 (0.0%) | |
| Pos (n=119) | 62 (84.9%) | 42 (89.4%) | 15 (100.0%) | |

[a]Tumor PD-L1 expression score as a maximum (max) score. PD-L1 expression was scored as negative (score=0), weak (score = 1), strong (score = 2) in all the tumor center/border TMA cores (2 x center; 2 x border for each patient tumor) and the highest score was selected as the patient-wise score in this analysis.

[b]EpiStain intensity was measured in Epi$^{+}$ cells in tumor center area (mean of two replicate TMA cores).

[c]Mesenchymal proteins (FAP, SPARC, VIM) were scored in tumor center and border areas and positivity in either core was considered as positive for each marker separately.

p-value, Pearson Chi-square (Exact two-sided).

**Table S6**: Tumor PD-L1 and FAP association with sarcomatoid ccRCC in full cohort.

| | Sarcomatoid | | |
|---|---|---|---|
| **Variable** | **No (n=390)** | **Yes (n=22)** | **p** |
| **CD45⁺ count (center)[a]** | | | 0.158 |
| Low (n=134) | 127 (33.7%) | 7 (31.8%) | |
| Med (n=134) | 130 (34.5%) | 4 (18.2) | |
| High (n=131) | 120 (31.8%) | 11 (50.0%) | |
| na (n=13) | | | |
| **PD-L1 (tumor max)[b]** | | | **<0.001** |
| Neg (n=291) | 287 (73.6%) | 4 (18.2%) | |
| Weak (n=97) | 85 (21.8%) | 12 (54.5%) | |
| Strong (n=24) | 18 (4.6%) | 6 (27.3%) | |
| **FAP (tumor max)** | | | **0.001** |
| Neg (n=280) | 269 (69.0%) | 11 (50.0%) | |
| Weak (n=88) | 85 (21.8%) | 3 (13.6%) | |
| Strong (n=44) | 36 (9.2%) | 8 (36.4%) | |
| **FAP⁺ stroma count[c]** | | | **<0.001** |
| Low (n=127) | 128 (35.3%) | 2 (9.5%) | |
| Medium (n=128) | 124 (35.2%) | 3 (14.3%) | |
| High (n=129) | 111 (30.6%) | 16 (76.2%) | |
| na (n=28) | | | |
| **EpiStain (center)** | | | 0.358 |
| Low (n=132) | 122 (32.4%) | 10 (45.5%) | |
| Medium (n=133) | 126 (33.4%) | 7 (31.8%) | |
| High (n=134) | 67 (34.2%) | 5 (22.7%) | |

[a]CD45$^+$ cell counts as fractions of Epi$^-$ cells analyzed in tumor center cores (mean of 2 replicates). The categorization into low, med, and high counts is based on the 3-tier equal categorization derived from the full ccRCC cohort categorization.

[b]Tumor PD-L1 and FAP expression scores as a maximum (max) score, as depicted in the Methods.

[c]FAP$^+$ stroma cell counts as fractions of Epi$^-$ cells analyzed in tumor border cores (mean of 2 replicates). The categorization into low, med, and high counts is based on the 3-tier equal categorization derived from the full ccRCC cohort categorization.

p-value, Pearson Chi-square (Exact two-sided).

**Table S7**: Tumor PD-L1 and FAP association with sarcomatoid ccRCC in patients with high tumoral leukocyte infiltration ($CD45^{high}$).

| | Sarcomatoid | | |
|---|---|---|---|
| **Variable** | **No (n=120)** | **Yes (n=11)** | **p** |
| **PD-L1 (tumor max)[a]** | | | **<0.001** |
| Neg (n=72) | 72 (60.0%) | 0 (0.0%) | |
| Weak (n=44) | 36 (30.0%) | 8 (72.7%) | |
| Strong (n=15) | 12 (10.0%) | 3 (27.3%) | |
| **FAP (tumor max)** | | | 0.113 |
| Neg (n=64) | 58 (48.3%) | 6 (54.5%) | |
| Weak (n=43) | 42 (35.0%) | 1 (9.1%) | |
| Strong (n=24) | 20 (16.7%) | 4 (36.4%) | |
| **$FAP^{+}$ stroma count[b]** | | | 0.311 |
| Low (n=27) | 25 (21.9%) | 2 (18.2%) | |
| Medium (n=33) | 32 (28.1%) | 1 (9.1%) | |
| High (n=65) | 57 (50.0%) | 8 (72.7%) | |
| **EpiStain (center)** | | | 0.856 |
| Low (n=46) | 42 (35.0%) | 4 (36.4%) | |
| Medium (n=51) | 46 (38.3%) | 5 (45.5%) | |
| High (n=34) | 32 (26.7%) | 2 (18.2%) | |

[a]Tumor PD-L1 and FAP expression scores as a maximum (max) score.

[b]$FAP^{+}$ stroma cell counts as fractions of $Epi^{-}$ cells analyzed in tumor border cores (mean of 2 replicates). The categorization into low, med, and high counts is based on the 3-tier equal categorization derived from the full ccRCC cohort categorization.

p-value, Pearson Chi-square (Exact two-sided).

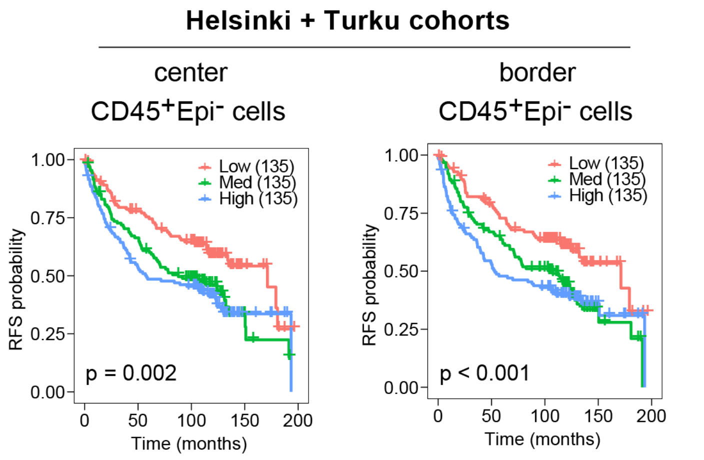


Cox regression continuous values: tumor center: $p = 0.004$, HR = 5.66

tumor border: $p < 0.001$, HR = 10.27

**Figure S1. Validation of CD45+ cell count association with worse RFS.**

Kaplan-Meier survival curves for recurrence-free survival (RFS) based on trichotomized proportions of $CD45^{+}Epi^{-}$ cells in different tumor center and border cores separately. Values as mean counts from replicate TMA cores. Macrophage multiplex panel. The three-tiered categorization was done after merging the Helsinki and Turku cohorts.

p-values from log-rank test.

Cox regression p-values and hazard ratios (HR) shown below the Kaplan-Meier curves indicate that the survival effects are significant also when analyzed using continuous values for cell counts.

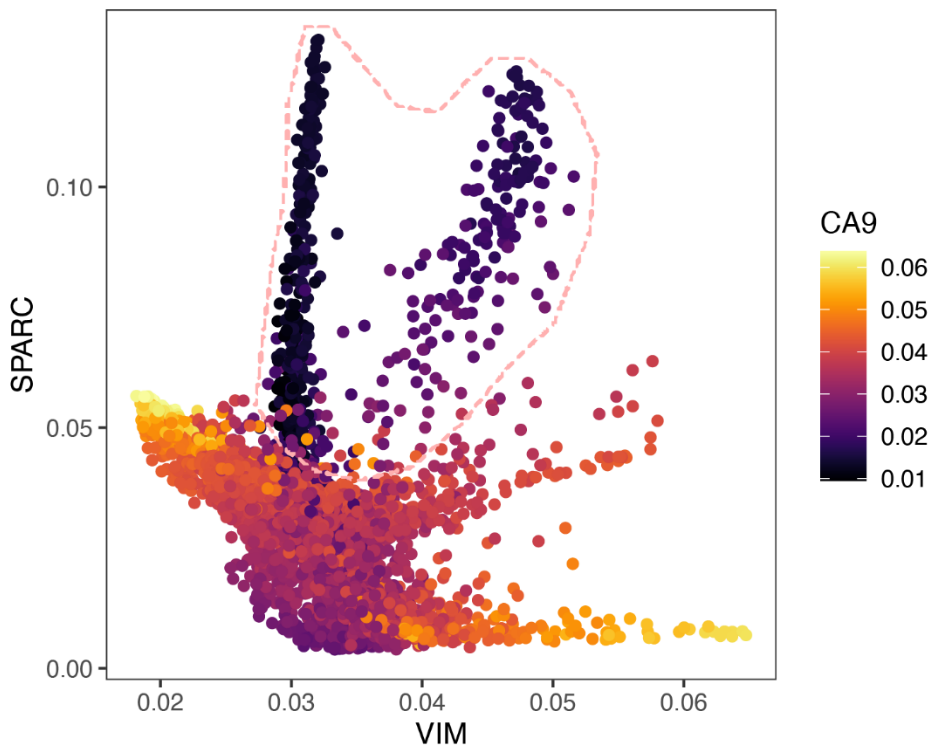


**Figure S2. Single-cell scatter plot of VIM and SPARC expression in RCC tumor cells with CA9 expression annotation**

Analysis of single-cell RNA-sequencing dataset from Li et al.[1] The analysis was performed on cells annotated as RCC by the authors of the original paper. The dotted line highlights clusters of tumor cells positive for mesenchymal markers VIM and SPARC but showing absent or very low expression signal for the CA9 epithelial marker. See Methods for analysis details.

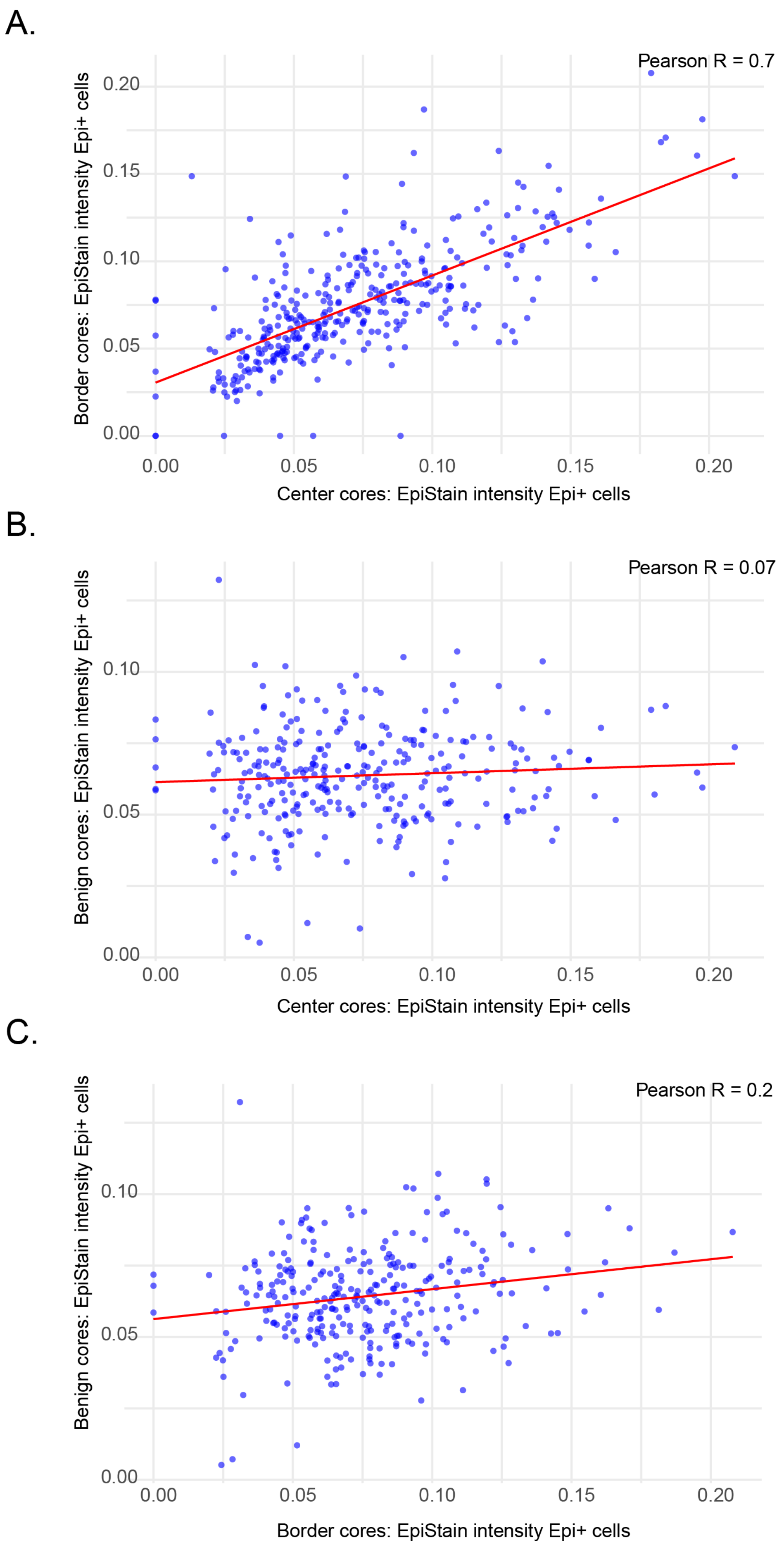
A.
Pearson R = 0.7
Border cores: EpiStain intensity Epi+ cells
0.00
0.05
0.10
0.15
0.20
0.00
0.05
0.10
0.15
0.20
Center cores: EpiStain intensity Epi+ cells
B.
Pearson R = 0.07
Benign cores: EpiStain intensity Epi+ cells
0.00
0.05
0.10
0.00
0.05
0.10
0.15
0.20
Center cores: EpiStain intensity Epi+ cells
C.
Pearson R = 0.2
Benign cores: EpiStain intensity Epi+ cells
0.00
0.05
0.10
0.00
0.05
0.10
0.15
0.20
Border cores: EpiStain intensity Epi+ cells

**Figure S3. Correlation plots of EpiStain intensity in Epi$^+$ cells between tumor center, border, and adjacent benign TMA cores.**

- **(A)** Scatter plot of tumor center and border Epi$^+$ cell EpiStain intensities (upper quartile intensity) as mean intensities per patient (two replicate cores per area). A linear regression trendline (red) is shown. The Pearson correlation coefficient (R) indicates a significant ($p < 0.001$) positive correlation between tumor center and tumor border EpiStain intensities across the patients. n = 357 patients.
- **(B)** Scatter plot of tumor center and benign Epi+ cell EpiStain intensities measured as in A. n = 301 patients.
- **(C)** Scatter plot of tumor border and benign Epi+ cell EpiStain intensities measured as in A. n = 290 patients.

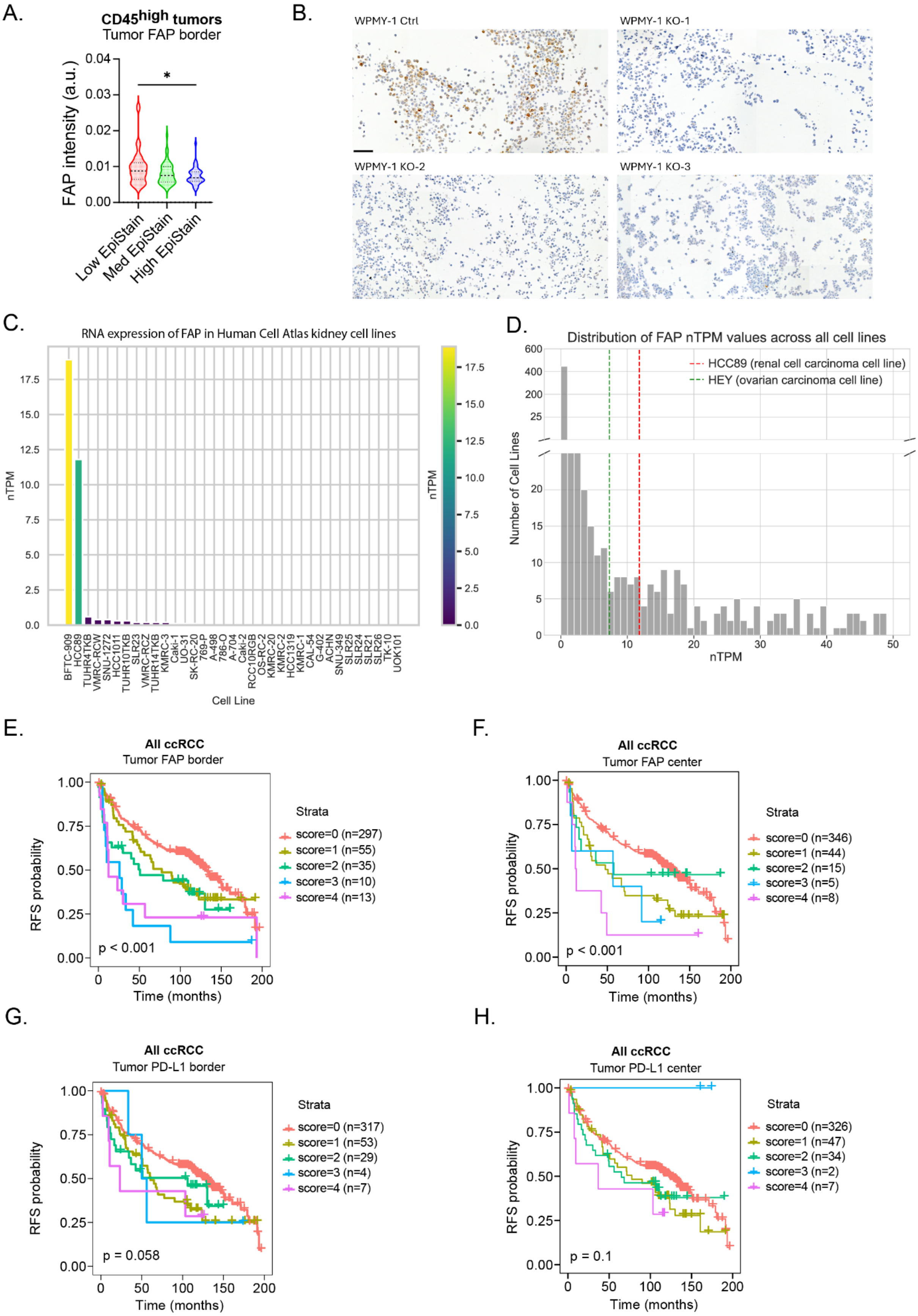
A.
CD45high tumors
Tumor FAP border
FAP intensity (a.u.)
Low EpiStain
Med EpiStain
High EpiStain
B.
WPMY-1 Ctrl
WPMY-1 KO-1
WPMY-1 KO-2
WPMY-1 KO-3
C.
RNA expression of FAP in Human Cell Atlas kidney cell lines
nTPM
Cell Line
D.
Distribution of FAP nTPM values across all cell lines
HCC89 (renal cell carcinoma cell line)
HEY (ovarian carcinoma cell line)
Number of Cell Lines
nTPM
E.
All ccRCC
Tumor FAP border
Strata
score=0 (n=297)
score=1 (n=55)
score=2 (n=35)
score=3 (n=10)
score=4 (n=13)
p < 0.001
RFS probability
Time (months)
F.
All ccRCC
Tumor FAP center
Strata
score=0 (n=346)
score=1 (n=44)
score=2 (n=15)
score=3 (n=5)
score=4 (n=8)
p < 0.001
G.
All ccRCC
Tumor PD-L1 border
Strata
score=0 (n=317)
score=1 (n=53)
score=2 (n=29)
score=3 (n=4)
score=4 (n=7)
p = 0.058
H.
All ccRCC
Tumor PD-L1 center
Strata
score=0 (n=326)
score=1 (n=47)
score=2 (n=34)
score=3 (n=2)
score=4 (n=7)
p = 0.1

**Figure S4. Association of tumor cell FAP and PD-L1 expression with recurrence-free survival (RFS) in tumor center cores of localized ccRCC.**

- **(A)** FAP expression (mean fluorescence intensity) in $Epi^+$ cells at the tumor border across low, medium, and high EpiStain intensity tumors (tumor center). *p < 0.05; Kruskal-Wallis test.
- **(B)** Validation of FAP antibody (ab207178) in IHC using WPMY-1 fibroblast cell line with CRISPR-CAS9 knock-out (KO) of control or *FAP* gene (three KO clones). Antibody dilution 1:500. Scale bar = 100 mm
- **(C)** RNA expression of FAP in kidney-derived cell lines in the Human Cell Atlas of The Human Protein Atlas version 24.0 (https://www.proteinatlas.org/humanproteome/cell+line/data#cell_lines). nTPM, normalized Transcripts Per Million.
- **(D)** Distribution of FAP nTPM values across cell lines highlighting the clear cell renal cell carcinoma cell line HCC89 (nTPM = 11) and ovarian carcinoma cell line HEY (nTPM = 7), which stands as a reference for a cell line with experimentally validated FAP expression.[2]
- **(E)** Kaplan-Meier curves for tumor cell-specific FAP expression in $CD45^{high}$ ccRCCs (n = 135) based on 3-tier visual scoring. FAP was scored (negative=0, weak=1, strong=2) in tumor center TMA cores (n = 2) and the scores were summed, yielding final scores from 0 to 4. See the cumulative scoring system in Method details.
- **(F)** Same as in B, but with the full ccRCC cohort (n=405).
- **(G)** Same as in B, but for tumor cell-specific PD-L1 expression (n=135).
- **(H)** Same as in C, but for tumor cell-specific PD-L1 expression (n=416).

p-values from log-rank test.

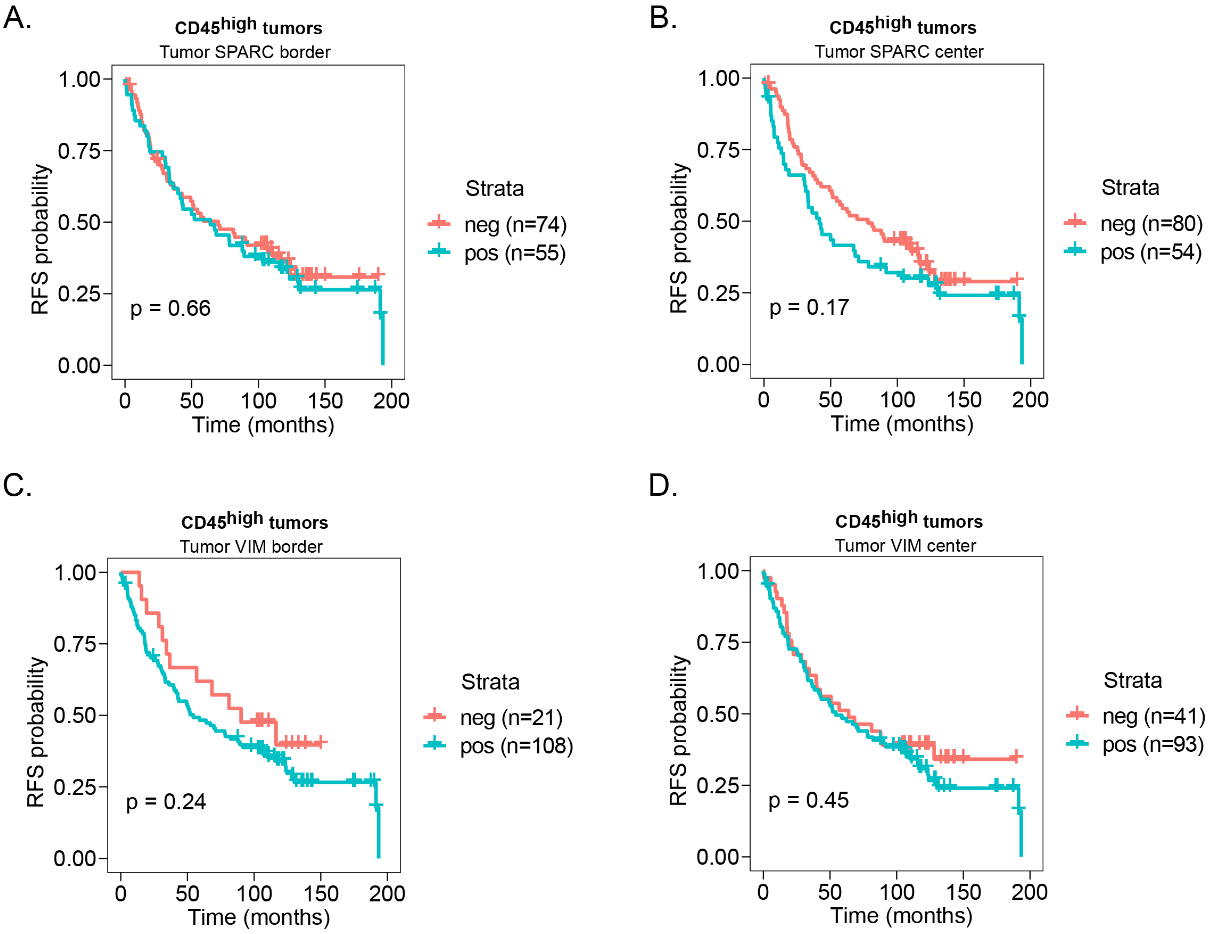


**Figure S5. Tumor cell SPARC and VIM association with RFS in localized CD45$^{high}$ ccRCC.**

- **(A)** Kaplan-Meier curves for tumor cell-specific SPARC expression in CD45$^{high}$ ccRCCs (n = 129) based on negative/postive scoring. SPARC was considered positive if either tumor border core (2 replicates) was positive.
- **(B)** Same as in A, but in tumor center cores (n = 134)
- **(C)** Kaplan-Meier curves for tumor cell-specific VIM expression in CD45$^{high}$ ccRCCs (n =129) based on negative/positive scoring. VIM was considered positive if either tumor border core (2 replicates) was positive.
- **(D)** Same as in C, but in tumor center cores (n = 134).

p-values from log-rank test.

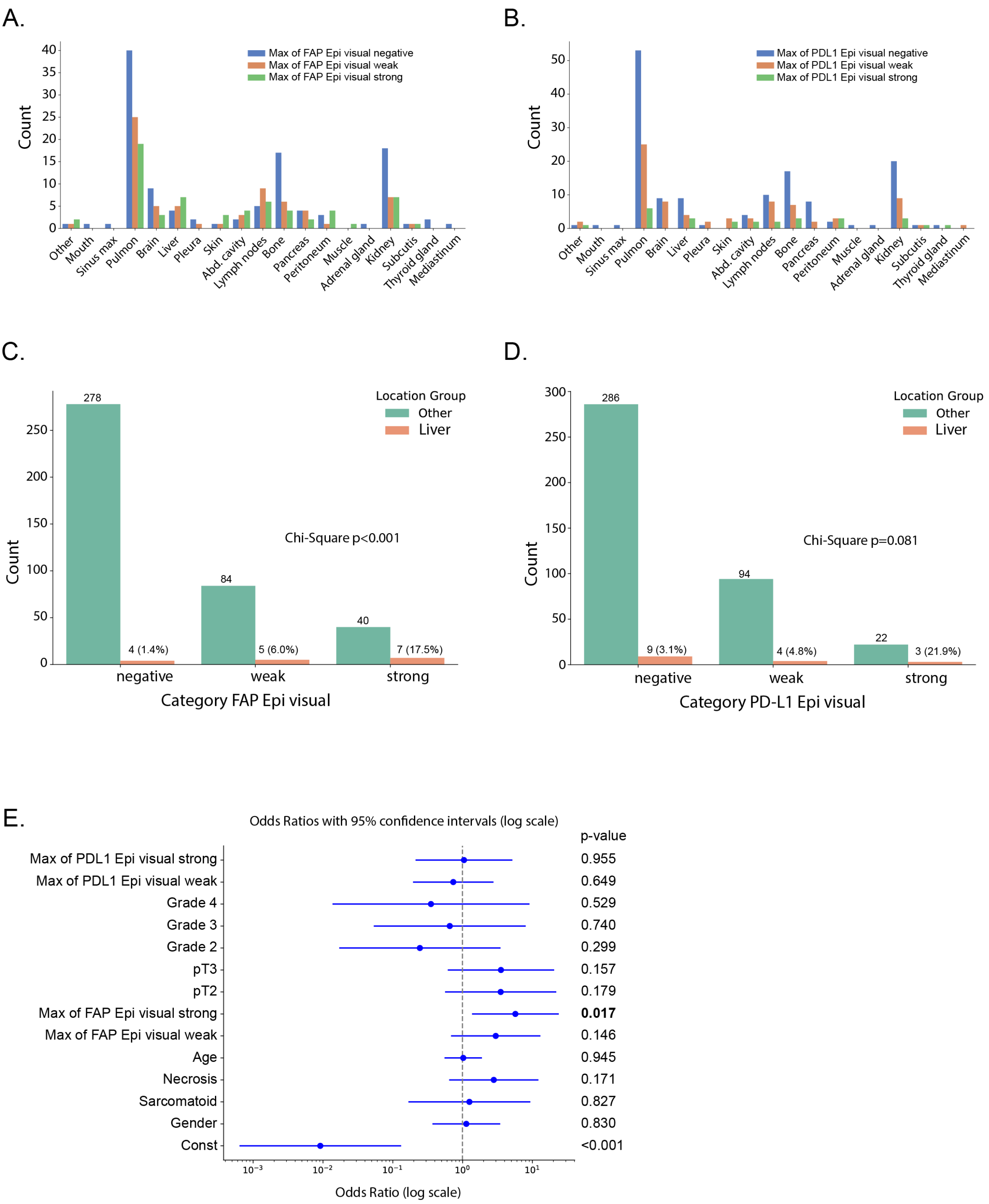
A.
Count
Max of FAP Epi visual negative
Max of FAP Epi visual weak
Max of FAP Epi visual strong
Other
Mouth
Sinus max
Pulmon
Brain
Liver
Pleura
Skin
Abd. cavity
Lymph nodes
Bone
Pancreas
Peritoneum
Muscle
Adrenal gland
Kidney
Subcutis
Thyroid gland
Mediastinum
B.
Count
Max of PDL1 Epi visual negative
Max of PDL1 Epi visual weak
Max of PDL1 Epi visual strong
C.
Location Group
Other
Liver
278
4 (1.4%)
84
5 (6.0%)
40
7 (17.5%)
Chi-Square p<0.001
Count
negative
weak
strong
Category FAP Epi visual
D.
Location Group
Other
Liver
286
9 (3.1%)
94
4 (4.8%)
22
3 (21.9%)
Chi-Square p=0.081
Count
negative
weak
strong
Category PD-L1 Epi visual
E.
Odds Ratios with 95% confidence intervals (log scale)
p-value
Max of PDL1 Epi visual strong 0.955
Max of PDL1 Epi visual weak 0.649
Grade 4 0.529
Grade 3 0.740
Grade 2 0.299
pT3 0.157
pT2 0.179
Max of FAP Epi visual strong 0.017
Max of FAP Epi visual weak 0.146
Age 0.945
Necrosis 0.171
Sarcomatoid 0.827
Gender 0.830
Const <0.001
Odds Ratio (log scale)

**Figure S6. Comparative analysis of FAP and PD-L1 expression in primary ccRCC tumors and their relationship with liver metastasis.**

- **(A)** Distribution of FAP expression across various metastasis locations identified through clinical PET-CT imaging. Categories of FAP expression in primary tumors are classified as negative, weak, and strong. Patients may have multiple metastasis sites; therefore, a single patient could be marked positive for several locations simultaneously, which explains the higher total count values across different metastasis locations.
- **(B)** Distribution of PD-L1 expression across different metastasis locations. Patients may have metastases in multiple locations, resulting in higher counts across locations.
- **(C)** Chi-square analysis comparing liver metastasis to other locations based on FAP expression levels (Chi-square $p < 0.001$).
- **(D)** Chi-square analysis comparing liver metastasis to other locations based on PD-L1 expression levels (Chi-square $p = 0.081$).
- **(E)** Logistic regression results showing Odds Ratios (OR) for FAP and PD-L1 expression categories as predictors of liver metastasis, controlling for clinical covariates. Strong FAP expression is a significant predictor of liver metastasis (OR = 5.70; $p = 0.017$), while PD-L1 expression shows no significant association with liver metastasis.

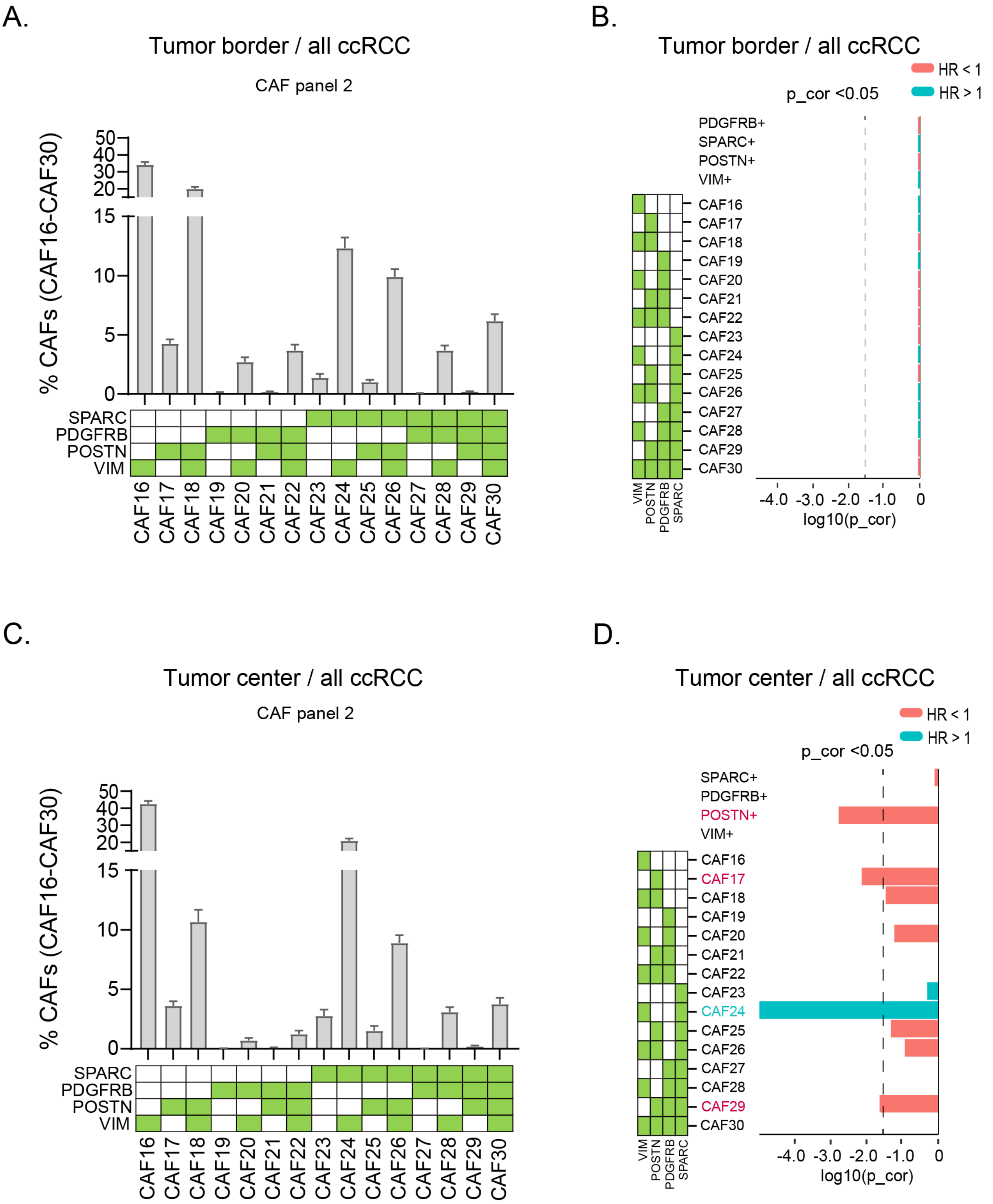
A.
Tumor border / all ccRCC
CAF panel 2
% CAFs (CAF16-CAF30)
50
40
30
20
15
10
5
0
SPARC
PDGFRB
POSTN
VIM
CAF16
CAF17
CAF18
CAF19
CAF20
CAF21
CAF22
CAF23
CAF24
CAF25
CAF26
CAF27
CAF28
CAF29
CAF30
B.
Tumor border / all ccRCC
HR < 1
HR > 1
p_cor <0.05
PDGFRB+
SPARC+
POSTN+
VIM+
CAF16
CAF17
CAF18
CAF19
CAF20
CAF21
CAF22
CAF23
CAF24
CAF25
CAF26
CAF27
CAF28
CAF29
CAF30
VIM
POSTN
PDGFRB
SPARC
-4.0 -3.0 -2.0 -1.0 0
log10(p_cor)
C.
Tumor center / all ccRCC
CAF panel 2
% CAFs (CAF16-CAF30)
50
40
30
20
15
10
5
0
SPARC
PDGFRB
POSTN
VIM
CAF16
CAF17
CAF18
CAF19
CAF20
CAF21
CAF22
CAF23
CAF24
CAF25
CAF26
CAF27
CAF28
CAF29
CAF30
D.
Tumor center / all ccRCC
HR < 1
HR > 1
p_cor <0.05
SPARC+
PDGFRB+
POSTN+
VIM+
CAF16
CAF17
CAF18
CAF19
CAF20
CAF21
CAF22
CAF23
CAF24
CAF25
CAF26
CAF27
CAF28
CAF29
CAF30
VIM
POSTN
PDGFRB
SPARC
-4.0 -3.0 -2.0 -1.0 0
log10(p_cor)

**Figure S7. Distribution and survival associations of CAFs in localized ccRCC using markers from CAF panel 2.**

- **(A)** Relative distributions of CAF panel 2 multi-marker defined stromal cell subsets in tumor border TMA cores (mean of replicates) (n = 408). Each CAF subset abundance is relative to the sum of the 15 CAFs as in [3]. Bars represent mean values and error bars 95% confidence interval.
- **(B)** Univariate Cox regression survival analysis using continuous values for the indicated cell subsets (all subsets $Epi^{-}$) in tumor border cores (mean of replicate cores).
- **(C)** Same as in A, but tumor center TMA cores for CAF panel 2 CAF subsets (n = 414).
- **(D)** Same as in B, but tumor center TMA cores for CAF panel 2 CAF subsets (n = 414).

Green box represents positivity. HR, Hazard ratio; p_cor, Bonferroni corrected p-value.

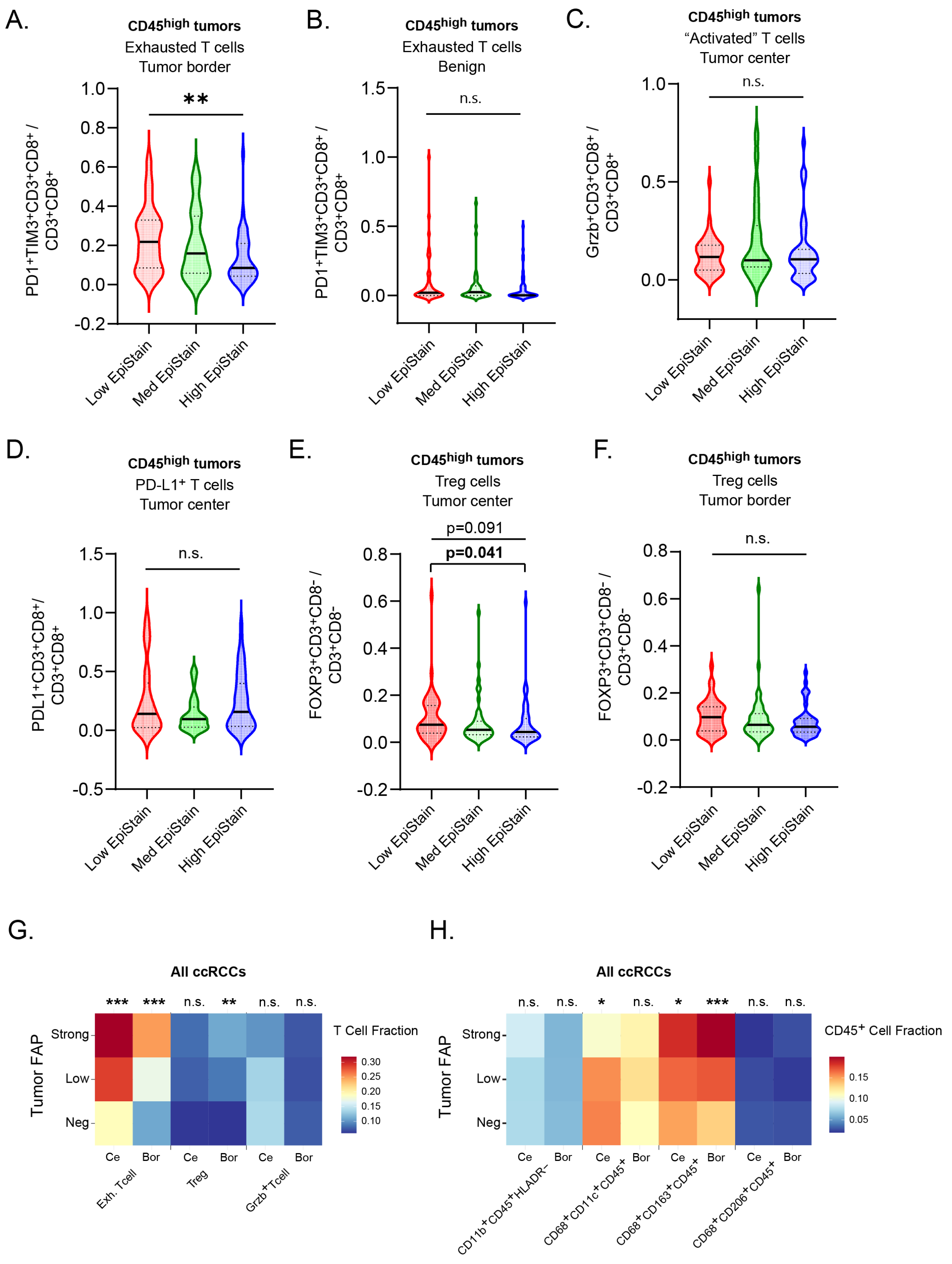
A.
CD45high tumors
Exhausted T cells
Tumor border
PD1+TIM3+CD3+CD8+ / CD3+CD8+
**
B.
CD45high tumors
Exhausted T cells
Benign
PD1+TIM3+CD3+CD8+ / CD3+CD8+
n.s.
C.
CD45high tumors
"Activated" T cells
Tumor center
Grzb+CD3+CD8+ / CD3+CD8+
n.s.
D.
CD45high tumors
PD-L1+ T cells
Tumor center
PDL1+CD3+CD8+/ CD3+CD8+
n.s.
E.
CD45high tumors
Treg cells
Tumor center
FOXP3+CD3+CD8- / CD3+CD8-
p=0.091
p=0.041
F.
CD45high tumors
Treg cells
Tumor border
FOXP3+CD3+CD8- / CD3+CD8-
n.s.
Low EpiStain
Med EpiStain
High EpiStain
G.
All ccRCCs
*** *** n.s. ** n.s. n.s.
Tumor FAP
Strong
Low
Neg
T Cell Fraction
Ce Bor Ce Bor Ce Bor
Exh. Tcell
Treg
Grzb+Tcell
H.
All ccRCCs
n.s. n.s. * n.s. * *** n.s. n.s.
CD45+ Cell Fraction
CD11b+CD45+HLADR-
CD68+CD11c+CD45+
CD68+CD163+CD45+
CD68+CD206+CD45+

**Figure S8. Distribution of immune cell subsets across EpiStain and tumor FAP categories in localized ccRCC.**

**(A)** Violin plots of distributions of $PD1^+TIM3^+CD3^+CD8^+$ cell subsets relative to total $CD3^+CD8^+$ T cells across low (n = 47), medium (Med, n = 49) and high EpiStain (n = 34) categorized tumors in tumor border TMA cores (mean of replicates) (total n = 130). Violin plot lines represent mean values and dotted lines inter quartile ranges (25 percentiles, 75 percentiles).

**(B)** Same as in A, but tumor adjacent benign TMA cores. n(low) = 41; n(med) = 47; n(high) = 31. (total n = 119).

**(C)** Distributions of $Grzb^+CD3^+CD8^+$ ('activated') cell subsets relative to total $CD3^+CD8^+$ T cells across low (n = 47), medium (Med, n = 51) and high EpiStain (n = 34) categorized tumors in tumor center TMA cores (mean of replicates) (total n = 132).

**(D)** Same as in C, but $PD\text{-}L1^+CD3^+CD8^+$ cell subsets. n(low) = 47; n(med) = 51; n(high) = 34. (total n = 132).

**(E)** Distributions of $FOXP3^+CD3^+CD8^-$ cell subsets relative to total $CD3^+CD8^-$ T cells across low (n = 48), medium (Med, n = 52) and high EpiStain (n = 35) categorized tumors in tumor center TMA cores (mean of replicates) (total n = 135). p-values indicate Kruskal-Wallis test between three variables (upper) and Mann-Whitney Exact two-sided test for comparing two variables (low vs. high).

**(F)** Same as in E, but tumor border TMA cores. n(low) = 47; n(med) = 51; n(high) = 34. (total n = 132).

**(G)** Heatmap of T cell fraction by cell type and tumor FAP level in full ccRCC cohort. Colors represent the relative fractions of effector T cells ($CD3^+CD8^+$) using a reversed RdYlBlu color palette. Tumor FAP scored as negative/weak/strong if either tumor center or border showed positivity (maximum level selected from center/border). T cell fractions as continuous values in tumor center (Ce) and border (Bor) TMA cores (average of replicates). n.s, not significant; **$p < 0.01$, ***$p < 0.001$; Kruskal-Wallis test.

**(H)** Same as G, but myeloid cell fractions of $CD45^+$ cells. n.s, not significant; *$p < 0.05$, **$p < 0.01$, ***$p < 0.001$; Kruskal-Wallis test.
